# Boltz-Perturb: Improving Diversity and Accuracy in Protein-Ligand Co-Folding through Training-Free Conditioning Perturbation

**DOI:** 10.64898/2026.08.05.742877

**Authors:** Hyeyun Jung, BoRam Lee, Alan C. Cheng

**Author notes:** Equal contribution (Hyeyun Jung, BoRam Lee). Correspondence to: BoRam Lee < >, Alan C. Cheng < >.

## Abstract

Protein-ligand co-folding models hold promise in structure-based drug discovery and small molecule interaction prediction, but often fail in predicting correct small molecule binding poses. We present Boltz-Perturb, a framework for addressing this through perturbing model conditioning signals during model inference, and show that such perturbations improve correct ligand binding mode predictions. We first show with true-coordinate injection experiments that the model’s learned energy landscape contains correct binding-mode basins, allowing us to reframe the problem as one of sampling deficiency. We then introduce two inference-time perturbation strategies, Token Bias Perturbation (TBP) and Token Conditioning Perturbation (TCP), which increase exploration of alternative binding poses. Across diverse protein–ligand systems, TCP improves top-20 oracle success rates by 2.6 to 7.8 fold. Boltz-Perturb attains higher oracle success rates compared to the Boltz-2 high diffusion temperature variant while requiring over 75% less compute. To our knowledge, this is the first systematic perturbation analysis of a co-folding architecture for small-molecule binding mode diversity. We demonstrate that inference-time perturbations can unlock latent structural diversity in generative co-folding models and improve protein-ligand predictions without costly retraining.

## 1. Introduction

Co-folding models such as AlphaFold 3 (AF3), RosettaFold-AA, and open-source AF3-style co-folding models such as Boltz, Chai, Protenix, SeedFold, and OpenFold3 have transformed all-atom biomolecular structure prediction, including for small-molecule complexes with proteins (Abramson et al., 2024; Wohlwend et al., 2024; Krishna et al., 2024; Protenix Team et al., 2026; The OpenFold3 Team, 2025; Passaro et al., 2025; Zhou et al., 2025). On multiple benchmarks, these models can be competitive with classical docking methods (McNutt et al., 2025) in protein-ligand binding pose prediction, performing especially well in situations with moderate to high training-set similarity and where conformational changes are involved in binding (Zheng et al., 2025; Singh et al., 2026; Škrinjar et al., 2025). Despite these advances, critical limitations persist that undermine their utility for protein–ligand prediction and structure-based drug discovery.

Co-folding models frequently produce low-diversity outputs that converge to similar, often suboptimal binding modes. This occurs even when inference is repeated across multiple random seeds or diffusion sampling temperatures. This behavior reduces the effectiveness of downstream computational pipelines where accurate binding modes are essential for structure-activity relationships (Thaler et al., 2025).

This challenge is not unique to biomolecular modeling. In image and language generation, analogous low-diversity issues in conditional diffusion models have been addressed through architectural refinement, model finetuning, attention-level guidance, and conditioning perturbation (Zhang et al., 2024; Ho & Salimans, 2022; Wu et al., 2026; Sadat et al., 2023; Ahn et al., 2025b; Ronneberger et al., 2015). Inspired by these findings, we hypothesized that the low-diversity behavior of co-folding models is partly attributable to a sampling limitation and that broader exploration during reverse diffusion can improve predictions without retraining or fine-tuning.

AF3-style co-folding models have a trunk that processes inputs such as sequence and evolutionary information into trunk representations and a denoising module that produces predicted structures by iteratively refining atomic coordinates from noise (Abramson et al., 2024; Jumper et al., 2021). The denoising module is conditioned on these trunk-derived representations—per-token single representations and residue-pair pairwise representations. These representations are computed once and remain fixed across all structure samples generated for a given target (Abramson et al., 2024; Wohlwend et al., 2024; Passaro et al., 2025). We hypothesized that this static conditioning acts as a bottleneck that limits diversity. Here, we present Boltz-Perturb, a training-free perturbation framework that injects time-annealed noise (i.e., noise whose magnitude decreases over denoising steps) into these conditioning signals during inference (Figure 1). First, we conducted true-coordinate injection experiments showing that Boltz recovers correct binding modes when guided toward the correct conformational region (Bennett et al., 2023). This result suggests that the learned energy landscape can contain correct basins corresponding to experimentally observed binding modes, but default trajectories converge to suboptimal modes before reaching them. Second, inspired by the Condition-Annealed Diffusion Sampler (CADS) (Sadat et al., 2023) work, we introduce two complementary perturbation strategies in Boltz-2: Token Conditioning Perturbation (TCP), which targets the single representations conditioning token embeddings, and Token Bias Perturbation (TBP), which targets the attention bias derived from pairwise representations.

**Figure 1.**
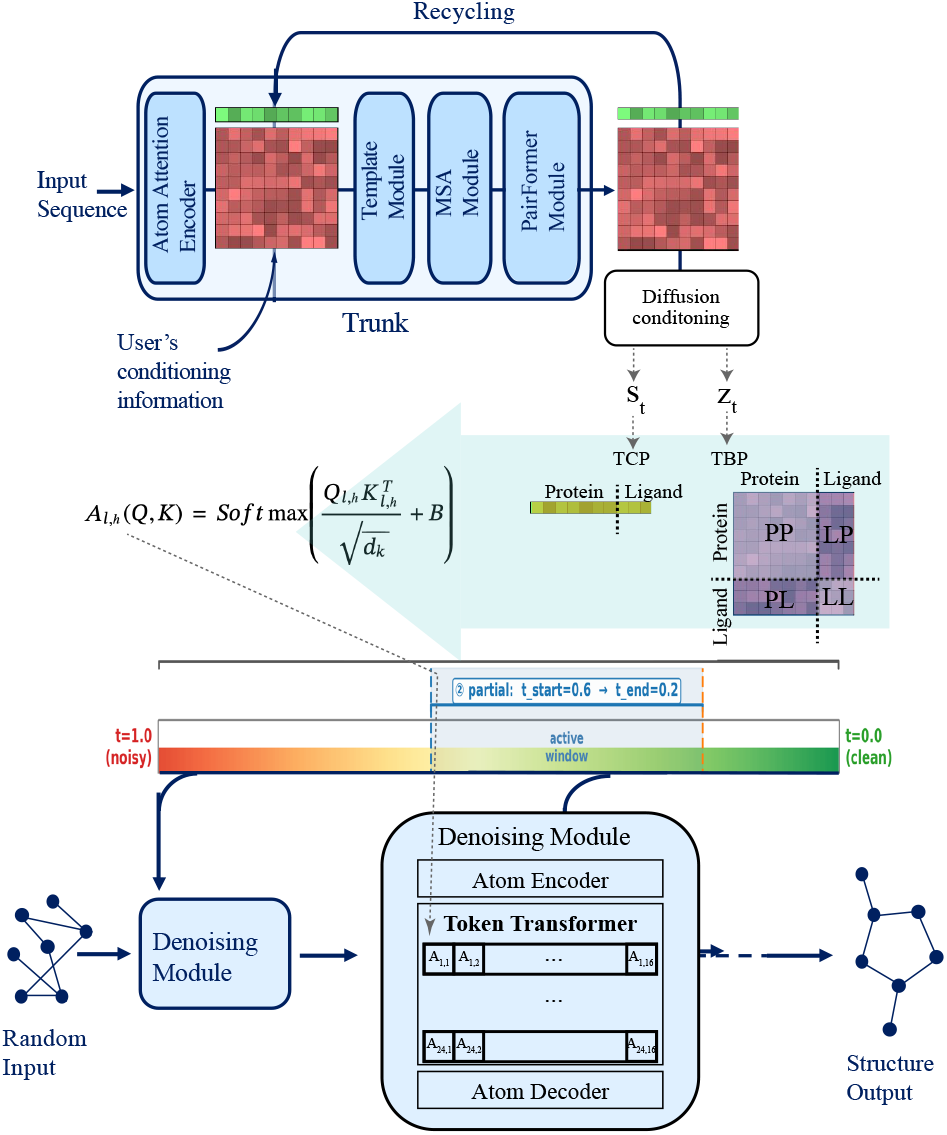
Overview of Boltz-Perturb.

Our main contributions are as follows:

1. **Diagnosing sampling deficiency**. Through true-coordinate injection experiments, we demonstrate that Boltz-2 possesses the latent capacity to identify correct binding modes, suggesting that the problem can be reframed as a sampling problem amenable to inference-time solutions.
2. **Boltz-Perturb**. We present a training-free framework that injects time-annealed noise at conditioning signals within the denoising module: token embeddings and attention biases. Without additional training, TCP achieved the highest rank in best ligand RMSD, improving the fraction of predictions with ligand RMSD lower than 2Å from 17.7% to 30.6% using three-fold fewer samples and overall lower additional compute cost.
3. **Characterizing the perturbation design space**. Through systematic empirical study, we characterize the relationship between noise scale, time-step scheduling, perturbation region to identify effective configurations. In addition, we introduce TADS-Auto, an adaptive schedule that uses the model’s own noise variance to automatically calibrate perturbation timing and magnitude.

## 2. Related work

### 2.1. Sampling Diversity for Diffusion Models

Despite the success of diffusion models in generating highquality outputs, they frequently suffer from low output diversity (Zhang et al., 2024). This lack of diversity is often amplified by strong conditioning guidance such as classifier-free guidance (CFG), which was developed to enhance sample quality by interpolating between conditional and unconditional score estimates during inference (Ho & Salimans, 2022; Dhariwal & Nichol, 2021). As the guidance scale increases, sample quality improves at the cost of diversity, with outputs clustering around dominant modes. Several training-free methods address this by perturbing the conditioning signal during inference. CADS introduced time-annealed conditioning perturbation (Sadat et al., 2023), corrupting the conditioning signal (*ŷ*) as:

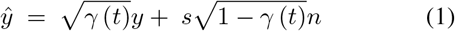

where *y* is the original conditioning signal, ***n*** is random Gaussian noise (*n* ~ *N* (0, 1)), *γ*(*t*) is a piecewise-linear annealing schedule, and *s* is an initial noise magnitude. The corrupted signal is then rescaled to preserve the original mean and variance, followed by interpolation for smoother control. CADS provides a principled justification through a score smoothing interpretation. It shows that the noise variance acts as ridge regularization on the score gradient that can reduce the dominance of any single mode and smoothes the conditional landscape.

Similarly, TAPS (Time-Annealed Perturbation Sampling) perturbs the conditioning signal during inference in diffusion language models (Wu et al., 2026). TADA (Training-free Augmented Dynamics) shows that pseudo-noise improves output diversity (Ahn et al., 2025b). These methods collectively establish that time-annealed conditioning perturbation is a principled, training-free mechanism for restoring diversity in diffusion models. Our work extends this principle to the biomolecular co-folding setting, where the conditioning signals are high-dimensional structured tensors, pairwise representations, and token embeddings.

### 2.2. Diversity Strategies in Structure Prediction

Several strategies have been proposed to increase conformational diversity in structure prediction, targeting different intervention points including temperature and seed variation, fine-tuning on additional structures, and altering inputs through MSA perturbation (del Alamo et al., 2022). MSA subsampling, random masking MSA, and clustering-based approaches perturb the input multiple sequence alignment to induce structural diversity, especially for fold-switching proteins (Kalakoti & Wallner, 2024).

Boltz-steering potentials apply physics-based potential gradient signals to the predicted denoising coordinates during reverse diffusion, biasing trajectories toward specific physical objectives. Recent concurrent work, ConforMix, employs twisted sequential Monte Carlo sampling with RMSD-based guidance potentials to bias predictions away from a reference structure, enabling undirected conformational exploration through particle filtering and resampling (Richman et al., 2025). For latent-based approaches, ConforNets requires additional training for lightweight channel-wise affine transforms on pre-Pairformer pair latents to modulate residue-residue contact processing, enabling both unsupervised diversity maximization and supervised conformational transfer. ConforMix and ConforNets focus on protein backbone diversity rather than ligand binding modes (Lee et al., 2026).

Our approach differs from existing methods in two ways. First, unlike MSA perturbation methods, we intervene directly in the latent representation of the denoising module, leaving the input pipeline unchanged. Second, while coordinate-level methods such as Boltz-steering and ConforMix optimize toward specific physical objectives or bias away from reference structures, we apply stochastic perturbation for broader conformational exploration without any predefined goal. Like coordinate-level approaches, our method is training-free and operates at inference time, making the two strategies complementary and combinable.

## 3. Methods

### 3.1. Boltz-2 Denoising Module Conditioning Signals

AF3-style co-folding models condition their denoising modules through trunk-derived representations. The trunk produces two primary conditioning signals, a single representation (*s*_*i*_) and a pairwise representation (*z*_*ij*_). In the denoising module’s token transformer, the single conditioning signal *s*_*i*_, which incorporates a Fourier embedding of the diffusion timestep *t*, modulates the token hidden state in each transformer block and is used to derive the query, key, and value projections. The pairwise conditioning signal *z*_*ij*_ is projected and integrated into the attention mechanism as an additive logit bias *B*:

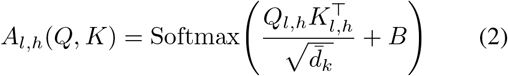

where *Q*_*l,h*_, *K*_*l,h*_ are query and key projections for the *l*-th layer and *h*-th head, and 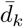 is the per-head dimensionality (Wohlwend et al., 2024; Passaro et al., 2025).

Because the trunk is computed only once, both conditioning signals are shared identically across all denoising samples. Although *s*_*i*_ varies across diffusion steps through its timestep embedding, it remains identical across samples at any given step. The bias *B* is fully static. Additionally, empirical analysis shows that the pairwise bias *B* strongly influences protein-side attention, leading to highly similar contact maps across samples. Recycling, if used, provides only subtle adjustments to the conditioning signals through reevaluating the trunk, and these updates are minor compared to the conformational diversity explored in the diffusion module.

In the following sections we introduce Token Conditioning Perturbation (TCP) and Token Bias Perturbation (TBP), which perturb the conditioning signals derived from *s*_*i*_ and *z*_*ij*_, respectively, in the token transformer, making each denoising sample unique and breaking this bottleneck. Each method consists of two components: (1) a noise schedule that determines the perturbation magnitude at each diffusion step and (2) a perturbation generator that constructs and applies sample-specific noise (Algorithms 1, 2, and S1). Additional details, including a full theoretical derivation, are provided in the Supplemental Information (Section A).

### 3.2. Perturbation Noise Scheduling (*λ* (*t*))

The diffusion sampling process alternates between adding noise and denoising. Therefore, any additionally injected perturbation should be aware of the current denoising step. If the perturbation is too strong, it risks overwhelming the signal that the model has already recovered. If it is too weak, the model may not respond to it at all. Thus, our perturbation uses time adaptive diffusion scaling (TADS) for noise scale (*λ*(*t*)) following CADS scheduling:

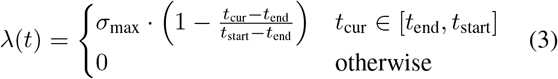

where *t*_cur_ = 1 − (*t/T*), *t* is the current denoising step, and *T* is the total number of denoising steps (Sadat et al., 2023). TADS schedules noise such that the perturbation noise *λ*(*t*) decays linearly within a given injection window [*t*_end_, *t*_start_] as a function of diffusion timestep *t* and given an initial noise level *σ*_max_.

We further introduce TADS-Auto for noise scheduling (Algorithm 1) to reduce reliance on manual hyperparameter tuning. TADS-Auto automatically determines the appropriate injection timing and scale by monitoring the model’s own noise during the diffusion process. Specifically, it monitors *ν*_*t*_, the standard deviation of the noise being added at step *t*, and activates perturbation only when *ν*_*t*_ is small enough that the injected noise would not be dominated by the model’s own stochastic noise. When *ν*_*t*_ ≤ *σ*_max_, two candidate scales are computed. The first, *λ*_1_(*t*) = *σ*_max_ ·*δ*, where *δ* = *ν*_*t*_*/*(*σ*_max_ +10^−8^), is a dynamic scale that reflects the ratio between current stochastic noise and the injected perturbation noise. The second, *λ*_2_(*t*) = *σ*_max_·*t*_cur_, is a steering scale that encodes a time-based linear decay proportional to diffusion progress. The final scale is *λ*_auto_ = max(*λ*_1_, *λ*_2_). This prevents the sudden zeroing of noise *λ* (*t*) = 0 when *ν*_*t*_ = 0 (i.e. no stochastic noise is being added), even if the diffusion has not ended. Therefore, TADS-Auto applies perturbation at the most sensitive diffusion timesteps automatically.

#### Algorithm 1

TADS-Auto

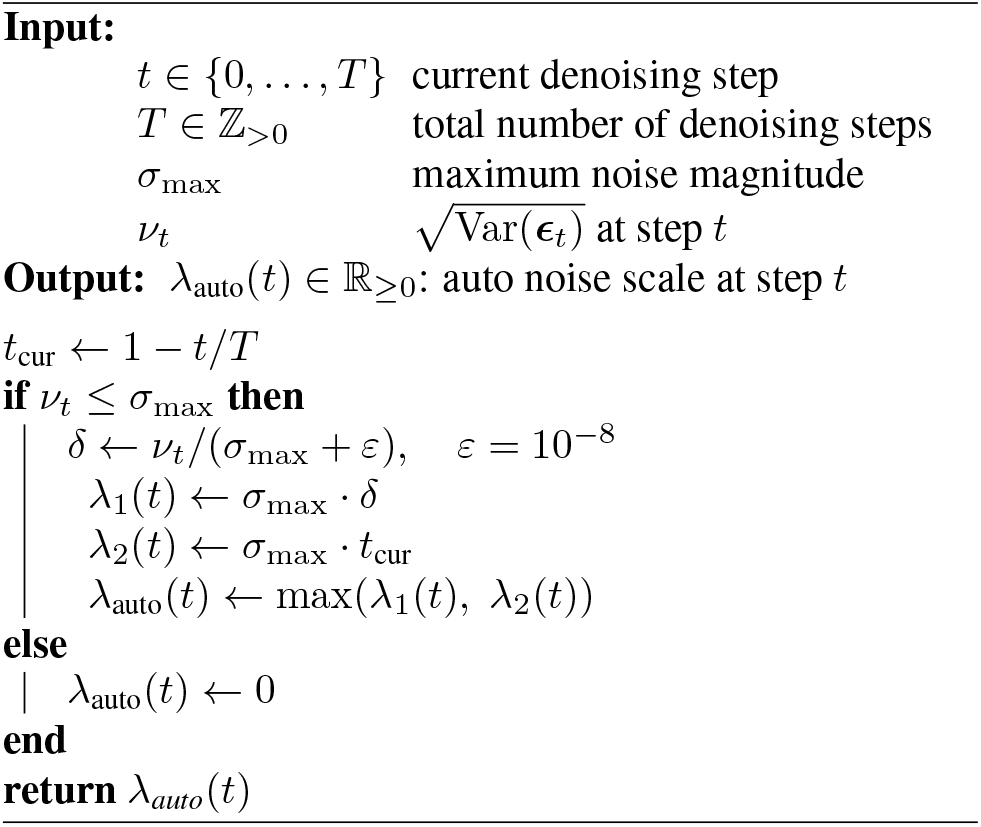

#### Algorithm 2

Boltz-Perturb: TCP/TBP

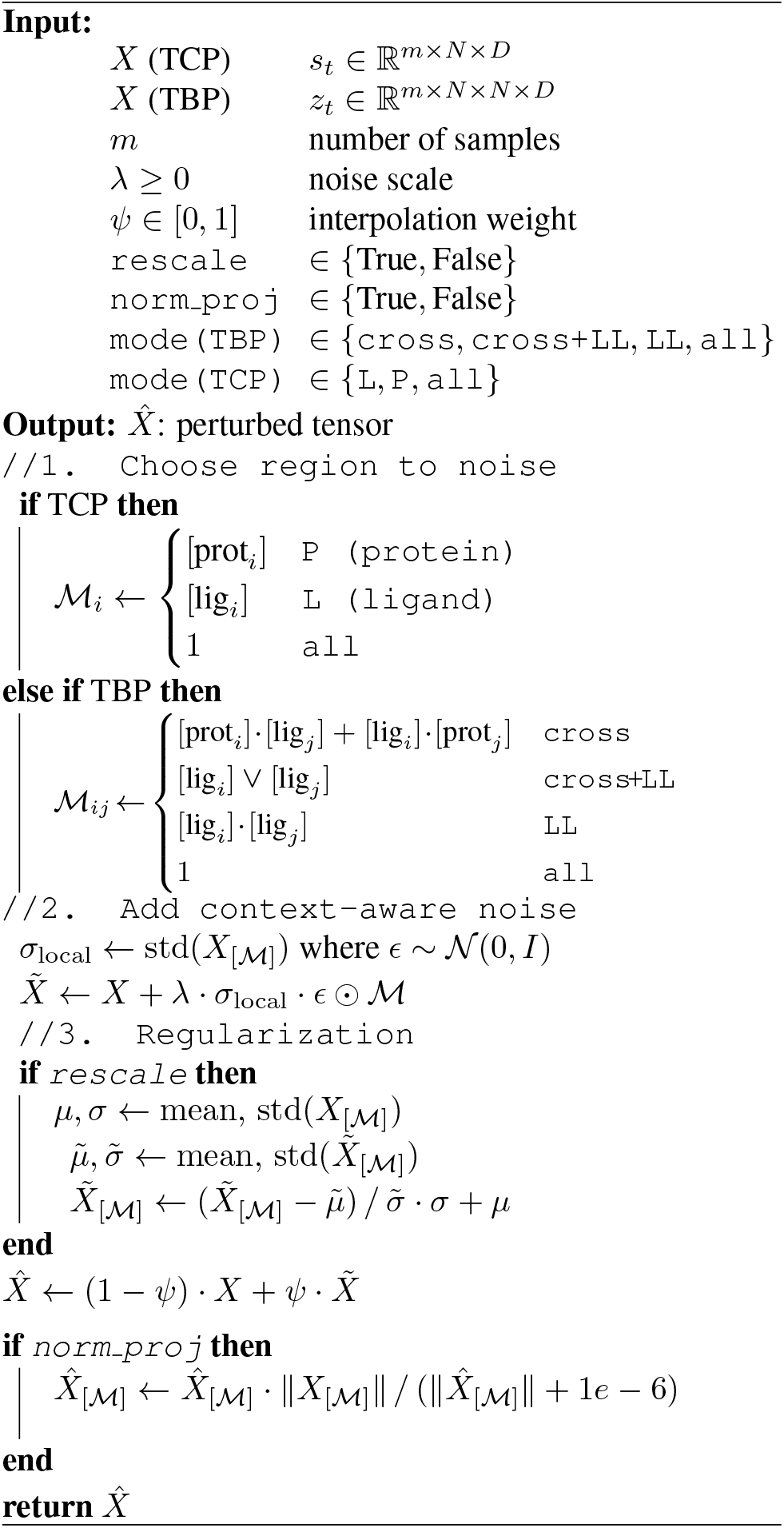

### 3.3. Perturbation Generation and Injection

After computing the noise scale *λ*(*t*), both TBP and TCP follow the same perturbation procedure but target different conditioning signals (Algorithm 2). TBP and TCP allow noise injection over a user-specified region defined by the molecule type. In this study, we focus on perturbing ligand tokens, as the ligand region is of more manageable size. To make the perturbation context-aware, we scale the injected noise by *λ*(*t*) *× σ*_local_, where *σ*_local_ is the standard deviation of the conditioning signal within the selected region *M*, calibrating the perturbation to intrinsic local variation rather than an arbitrary absolute scale. After the noise injection, we apply an interpolation factor (*ψ*) to linearly blend the original and perturbed signals to ensure smooth incorporation. We further implement two optional regularization strategies: *moment-matching rescaling*, which re-standardizes the perturbed signal to match the original mean and variance to prevent distributional drift (Sadat et al., 2023); and *norm-preserving projection*, which projects the perturbed signal onto the norm shell of the original, altering only its direction while preserving its magnitude (Wu et al., 2026; Sadat et al., 2023). Detailed perturbation conditions are provided in Table S1.

## 4 Experiments

### 4.1. Preliminary Diagnostic Experiment

Diffusion-based co-folding models encode richer conformational landscapes than their default sampling reveals (Richman et al., 2025). We evaluated this hypothesis on five recent PDB complexes absent from the Boltz-2 training data set. We generated 180 predictions per PDB complex using three random seeds and two temperatures. Vanilla Boltz-2 sampling produced a low diversity of structures, with ligand RMSF remaining below 2.3 Å in all cases and below 0.5 Å in two cases (Figures S1 and S2). These diagnostic results motivated us to focus on conditioning perturbation, a less invasive alternative to direct coordinate manipulation.

To test whether changing sampling in Boltz-2 alone can recover additional binding modes, we performed a coordinate injection experiment (additional details in Supplemental Information, Section B). At an intermediate denoising step, we replaced the predicted coordinates with the experimental ground truth a single time, and then continued standard reverse diffusion to completion. The final poses closely matched the reference structure, suggesting that the denoiser can refine to the correct binding mode once the trajectory enters its basin (Figure S1). However, because of high noise in the early diffusion schedule, coordinate injection at earlier steps reverted to the default result. Therefore, experimentally correct binding modes may be encoded within the model but remain underexplored by default sampling. Since the limitation is partially a sampling issue, we employed conditioning signal perturbation to redirect diffusion trajectories toward alternative conformational regions.

### 4.2. Experiment Set Up

#### Benchmarks

We evaluated on two sets: (1) a diagnostic set of five recently deposited protein–ligand complexes, and (2) 57 targets from the Runs N’ Poses (RnP) benchmark (Škrinjar et al., 2025). The RnP benchmark set was filtered to single-chain, single-ligand complexes deposited after the Boltz-2 training cutoff where the reported best vanilla prediction exceeded 2 Å ligand RMSD (details in Supplemental Information, sections B.2 and C).

#### Perturbation conditions

We evaluated two trunk-derived perturbation targets :

- **Token Bias Perturbation (TBP):** injects noise into the attention bias of the token transformer, which is derived from the pairwise representation *z*_*ij*_ and processed through diffusion conditioning. We vary perturbation region (`cross`, `cross+LL`, `LL`), noise schedule (TADS with fixed windows, TADS-Auto), and regularization (moment-matching rescaling, norm projection, interpolation *ψ*).
- **Token Conditioning Perturbation (TCP):** injects noise into the single conditioning signal *s*_*t*_ after diffusion conditioning, with perturbation restricted to ligand tokens for this study. TCP shares the same scheduling and regularization variants as TBP.

The full hyperparameter grid, including schedule windows and regularization ablations, is given in Supplemental Information (Section D.1).

#### Baselines

We compared TBP and TCP against five diversity strategies: (1) vanilla Boltz-2 (180 samples), (2) steering potentials, (3) elevated diffusion temperature (*T* ∈ {1.2, 1.3, 1.4}; total 270 samples), (4) MSA masking, and (5) MSA subsampling. These span physical guidance, temperature scaling, and input perturbation approaches (Supplemental Information, section B.3).

#### Sampling budget

Each perturbation condition generated 60 samples per target, a three-fold reduction relative to the vanilla budget of 180 samples and a 78% reduction relative to the vanilla high-temperature sampling budget of 270 samples.

#### Metrics

We evaluated ligand RMSD to ground truth, success rate (fraction of targets with ≥ 1 pose below 2 Å RMSD), model confidence, ligand RMSF with and without protein context, structural validity via PoseBusters. Metric definitions are in the Supplemental Information (Section C).

#### Compute

Experiments were run primarily on H100 (80 GB), NVIDIA A100 (80 GB), or V100 (32 GB) GPUs depending on availability. We observed GPU-dependent numerical variability under perturbation, which is a known issue (details in Supplemental Information, Section D.5).

## 5. Results

### 5.1. Boltz-Perturb improved accuracy and diversity

#### Diagnostic Five

Across all five targets, at least one perturbation condition achieved a ligand RMSD below 2 (Table 1 and Figure 2; full results in Table S2 and Figure S3).

**Table 1.** Minimum ligand RMSD (Å) per structure across baseline and perturbation methods. **Bold** = best, <u>underline</u> = second best. Abbreviations and condition details are defined below.^*†*^ For more and detailed results see Supplemental Information (Section D.3).

| PDB | Baselines |  |  |  |  | TBP |  |  |  | TCP |  |  |
| --- | --- | --- | --- | --- | --- | --- | --- | --- | --- | --- | --- | --- |
|  | V | V<sub>x</sub> | V<sub>hT</sub> | V<sub>mask</sub> | V<sub>sub</sub> | C1 | C2 | C3 | C4 | C10 | C11 | C12 |
| 9JF4 | 2.451 | 2.247 | 2.457 | 2.866 | 2.338 | 2.533 | 2.374 | <u>2.111</u> | 2.501 | 2.232 | **1.757** | 2.368 |
| 9M4Q | 2.433 | 3.014 | 3.015 | 3.019 | 3.061 | **1.897** | 2.902 | <u>1.921</u> | 2.372 | 2.670 | 2.325 | 2.918 |
| 9PY4 | 8.958 | 9.009 | 8.947 | 8.975 | 9.016 | 8.695 | **1.674** | 8.795 | 2.032 | <u>1.858</u> | 1.959 | 9.020 |
| 9RAY | 2.089 | 3.815 | 2.158 | 1.986 | 3.997 | 2.036 | 3.238 | 2.238 | **1.055** | 2.060 | <u>1.990</u> | 5.467 |
| 9Z1L | 7.212 | 7.205 | 7.195 | 7.214 | 7.213 | 6.874 | 1.154 | 6.885 | 1.073 | **0.968** | <u>1.068</u> | 7.180 |

**Baselines.** V: Boltz-2 vanilla (default); V<sub>x</sub>: Boltz-2x (steering); V<sub>hT</sub>: higher diffusion temperatures (1.2, 1.3, 1.4) vs. default (1.5, 1.638);
V<sub>m</sub>: MSA masking; V<sub>s</sub>: MSA subsampling.
**Perturbation methods.** TBP: Token Bias Perturbation (applied to protein–ligand token pairs); TCP: Token Conditioning Perturbation (applied to ligand tokens only). All TBP/TCP methods use normalization, rescaling, and interpolation factor $\psi=0.7$ . Configuration codes — C1: PL+LP+LL, $t_{\in[0.0,1.0]}$ ; C2: PL+LP, $t_{\in[0.2,0.6]}$ ; C3: PP+PL+LP+LL, $t_{\in[0.0,1.0]}$ ; C4: PL+LP, $t_{\text{auto}}$ ; C10: L, $t_{\in[0.0,1.0]}$ ; C11: L, $t_{\in[0.2,0.6]}$ ; C12: L, $t_{\text{auto}}$ .

**Figure 2.**
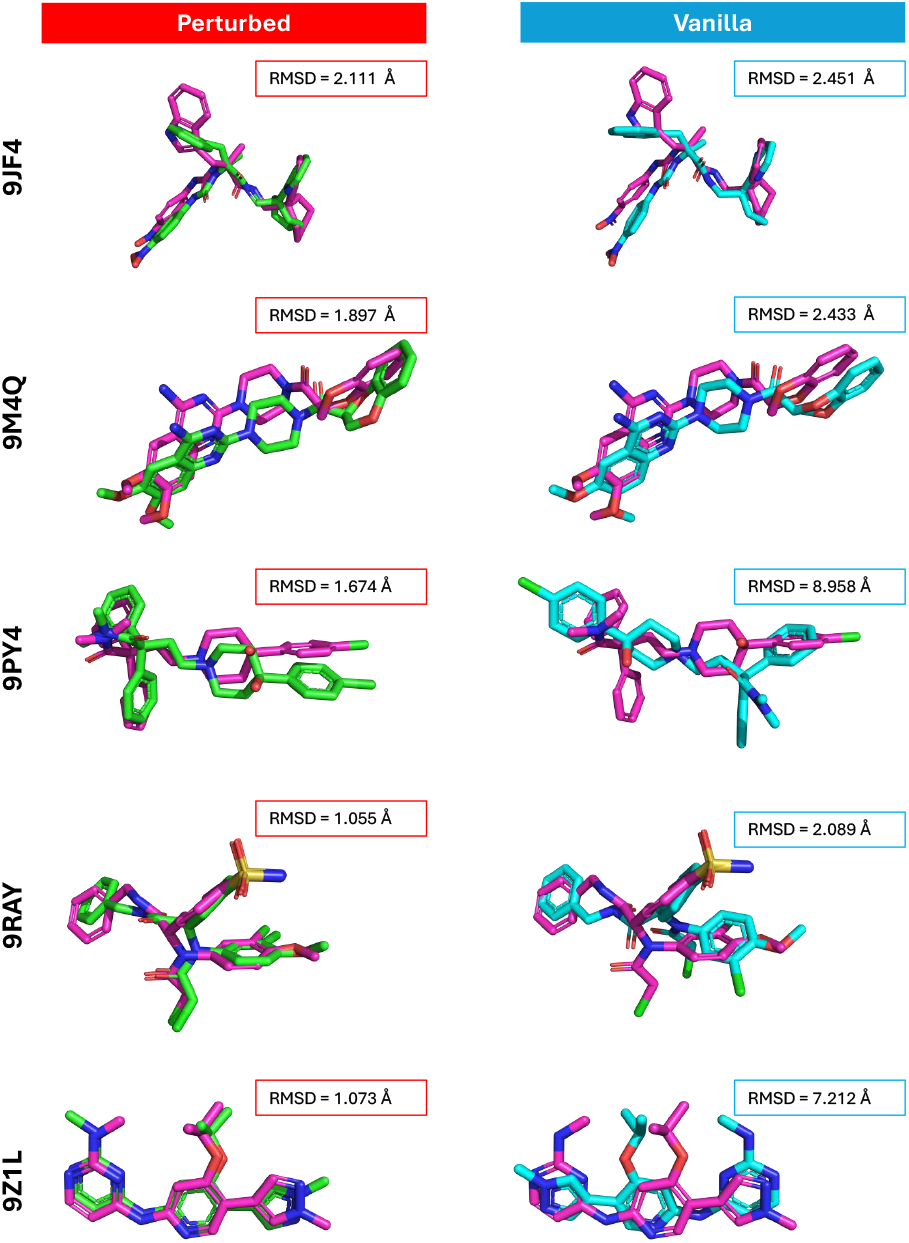
Comparison of the lowest RMSD structures identified in Table1. Left (TBP-perturbed, green), Right (Vanilla, blue). The ground truth experimental conformer is shown in magenta

The largest improvements were on 9PY4 and 9Z1L, where vanilla RMSD exceeded 7 Å (Table 1). For 9PY4, the best result came from mid-trajectory TBP perturbation of the protein-ligand cross-attention region with full regularization (C2), followed by TCP (C10). For 9Z1L, TCP with full-trajectory noise injection (C10) achieved the lowest RMSD, followed closely by mid-trajectory TCP (C11). For 9RAY, TBP was most effective when perturbation was restricted to the protein-ligand interaction region with auto-scheduled noise (C4).

#### RnP benchmark

Our vanilla Boltz-2 achieved a higher success rate than originally reported in RnP, likely due to the increased number of diffusion samples. Elevated temperature (V_hT_) improved oracle success rate but did not outperform our perturbation methods despite using 4.5-fold more samples. Other vanilla variations such as MSA masking, subsampling, and the Boltz steering potential did not improve RMSD accuracy. TCP-C11 achieved the lowest minimum RMSD on 20 of 57 targets (W_all_), far exceeding all other methods (Table 2).

**Table 2.** Comparison of perturbation methods across 57 targets from the Runs and Poses benchmark. (Predictions are all with NVIDIA H100 80 GB GPUs) **Bold** = best, <u>underline</u> = second best.

| Method | $N$ | Success Rate($<2 \text{ \AA}$) | | SR Oracle | | | Win | | Ligand RMSD | | RMSF ($\text{\AA}$)↑ | | |
| --- | --- | --- | --- | --- | --- | --- | --- | --- | --- | --- | --- | --- | --- |
| | | $SR_{20}(\%)$ | $SR_O(\%)$ | Ret | R↑ | D↓ | $W_{<2}$ | $W_{all}$ | $\bar{d} \pm \sigma \downarrow$ | | $F_p \pm \sigma$ | $F_l \pm \sigma$ | Val (%) |
| V | 180 | 5.26 | 19.30 | 11 | — | — | 3 | 3 | $9.604 \pm 7.025$ | | $2.382 \pm 2.613$ | $0.825 \pm 0.419$ | 98.79 |
| V<sub>x</sub> | 180 | 5.26 | 15.79 | 8 | 1 | 3 | 1 | 3 | $9.548 \pm 7.009$ | | $1.849 \pm 1.819$ | $0.724 \pm 0.378$ | 98.50 |
| V<sub>hT</sub> | 270 | 3.51 | <u>22.81</u> | **11** | 2 | **0** | 1 | 1 | $9.601 \pm 7.010$ | | $2.368 \pm 2.647$ | $0.802 \pm 0.389$ | 98.63 |
| V<sub>mask</sub> | 60 | 1.75 | 10.53 | 4 | 2 | 7 | 0 | 1 | $9.696 \pm 7.050$ | | $2.269 \pm 2.621$ | $0.793 \pm 0.378$ | 98.17 |
| V<sub>sub</sub> | 60 | 3.51 | 12.28 | 5 | 2 | 6 | 0 | 0 | $9.770 \pm 7.191$ | | $2.378 \pm 2.924$ | $0.795 \pm 0.376$ | 97.85 |
| TBP-C1 ($s=3.0$) | 60 | <u>10.53</u> | 15.79 | 6 | 3 | 5 | 1 | <u>7</u> | **9.324** $\pm 6.917$ | | $2.110 \pm 2.040$ | $0.869 \pm 0.389$ | 90.80 |
| TBP-C4 ($s=3.0$) | 60 | <u>10.53</u> | 14.04 | 6 | 2 | 5 | 1 | 1 | <u>9.467</u> $\pm 6.986$ | | $2.312 \pm 2.472$ | $0.807 \pm 0.376$ | 98.30 |
| TBP-C4 ($s=12.0$) | 60 | 7.02 | 17.54 | 7 | 3 | 4 | 2 | 4 | $9.725 \pm 7.005$ | | $2.768 \pm 2.260$ | $0.905 \pm 0.351$ | 98.48 |
| TBP-C4 ($s=25.0$) | 60 | 8.77 | 15.79 | 6 | 3 | 5 | <u>4</u> | 6 | $9.824 \pm 6.965$ | | $3.213 \pm 2.678$ | $0.922 \pm 0.381$ | 98.45 |
| TBP-C5 ($s=3.0$) | 60 | 7.02 | 14.04 | 7 | 1 | 4 | 0 | 2 | $9.896 \pm 7.195$ | | $2.768 \pm 3.382$ | $0.802 \pm 0.382$ | 98.80 |
| TBP-C5 ($s=12.0$) | 60 | 7.02 | <u>22.81</u> | 7 | 6 | 4 | 1 | 3 | $10.048 \pm 7.102$ | | $3.470 \pm 3.312$ | $0.908 \pm 0.387$ | 98.04 |
| TBP-C5 ($s=25.0$) | 60 | 7.02 | 17.54 | 5 | 5 | 6 | 2 | 4 | $10.166 \pm 7.114$ | | $3.892 \pm 3.682$ | <u>0.938</u> $\pm 0.390$ | 98.13 |
| TBP-C6 ($s=3.0$) | 60 | 7.02 | 12.28 | 6 | 1 | 5 | 1 | 2 | $9.467 \pm 7.039$ | | $2.056 \pm 2.176$ | $0.811 \pm 0.406$ | 98.52 |
| **TCP-C11 ($s=0.9$)** | **60** | **14.04** | **26.32** | <u>9</u> | **6** | <u>2</u> | **6** | **20** | $10.457 \pm 6.575$ | | **7.666** $\pm 4.241$ | **1.126** $\pm 0.485$ | 97.16 |

**Notation.** $N$ : total poses per target. V and V<sub>x</sub>: $3 \times 2 \times 30 = 180$ poses; V<sub>hT</sub>: $3 \times 3 \times 30 = 270$ poses; all other methods: $3 \times 2 \times 10 = 60$ poses. $SR_{20}$ : success rate (%) using top-20 confidence-ranked poses (success ⇔ at least one pose $< 2 \text{ \AA}$ per target). $SR_O$ : oracle success rate (best-case over all poses per target). Ret/R/D: number of targets retained/rescued/degraded relative to V\* at the $2 \text{ \AA}$ threshold (baseline: 11/57 successes). $W_{<2} / W_{all}$ : win count — number of targets (out of 57) where this method achieves the lowest min-RMSD across all methods, restricted to RMSD $< 2 \text{ \AA}$ ( $W_{<2}$ ) or unrestricted ( $W_{all}$ ). $\bar{d} \pm \sigma$ : mean ± std of per-target minimum ligand RMSD ( $\text{\AA}$ ) across 57 targets (lower is better). $F_p / F_l$ : mean ± std of per-target RMSF ( $\text{\AA}$ ) computed with/without protein context (higher indicates greater conformational diversity). Val: overall validity (valid poses/total poses).
**Method abbreviations.** TBP: Token Bias Perturbation; TCP: Token Conditioning Perturbation; V: Boltz-2 vanilla (default); V<sub>x</sub>: Boltz-2x (steering); V<sub>hT</sub>: higher diffusion temperatures (1.2, 1.3, 1.4) vs. default (1.5, 1.638); V<sub>m</sub>: MSA masking; V<sub>s</sub>: MSA subsampling. Configuration codes — C1: PL+LP+LL, $t_{\in[0.0,1.0]}$ ; C4: PL+LP, $t_{auto}$ ; C5: PL+LP, $t_{\in[0.0,1.0]}$ ; C6: PL+LP+LL, $t_{\in[0.2,0.6]}$ ; C11: L, $t_{\in[0.2,0.6]}$ (TCP variant). All TBP/TCP methods apply normalization, rescaling, and interpolation factor $\psi=0.7$ .

To assess whether the improvement was generalizable, we stratified success counts by SuCOS-pocket score (Figure 3), where lower scores indicate less similar pockets and ligands compared to training-set structures (Škrinjar et al., 2025). Vanilla Boltz-2 achieved 11 successes out of the 57 targets, primarily in pockets with mid-to-high similarity. TBP-C5 (*s*=12) improved the success rate with 13 successes and gains primarily in mid-to-high similarity regimes. TCP-C11 (*s*=0.9) improved it to 15 successes, with gains in mid-low and mid-high similarity regimes. Combining the unique successes of both TBP and TCP methods further increased the success rate to 21 out of 57 (Figure 3, SR Oracle column from Table 2). Although the lowest similarity bin (0–20, *N* =9) in Figure 3 contains too few targets to draw confident conclusions, we observed no improvement in this bin from TBP/TCP perturbations. This may imply that perturbation offer the greatest benefits in the mid-low and mid-high SuCOS-pocket similarity regimes. For targets with very low similarity to the training set, improving sampling alone may not sufficient and additional training may be necessary. Evaluation on an even broader set of diverse targets remains a key area for future work.

**Figure 3.**
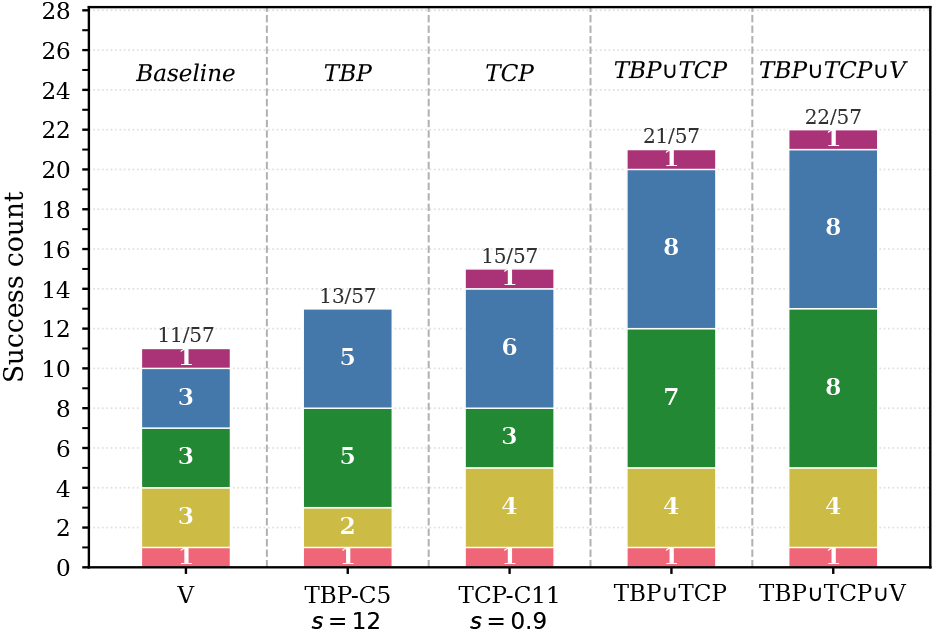
Success count (RMSD *<* 2 Å) for each condition (*x*-axis) for each PDB. If there is at least one pose that is *<* 2 Å for PDB A, then it is considered a success for PDB A. stratified by ligand pocket shape similarity (`sucos_shape_pocket_qcov_2023`). Bars are stacked by SUCOS bin: 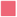: [0, 20) (*N* = 9), 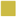: [20, 40) (*N* = 16), 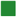: [40, 60) (*N* = 19), 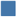: [60, 80) (*N* = 10), 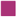:[80, 100) (*N* = 3). Numbers inside segments = per-bin count; labels above bars = total successes / *N* =57. Duplicates in TBP∪TCP and TBP∪TCP∪V counted once (V = vanilla Boltz-2).

The perturbation methods (TBP and TCP) outperformed all baselines in oracle success rate using only one-third of the sampling budget. Across the combined 62-target evaluation, TCP nearly doubled the oracle success rate from 17.7% to 30.6%. When considering only the top-20 confidence-ranked predictions per target, TBP and TCP showed substantially greater improvement over baselines (Figure 4; see also Table 2). To verify that our accuracy improvements stemmed from increased sampling diversity, we measured ligand RMSF across the generated samples. The best-performing conditions consistently showed higher RMSF than the baselines, both with and without protein context (Tables 1, 2) supporting our hypothesis that the perturbation methods drive broader conformational space exploration.

**Figure 4.**
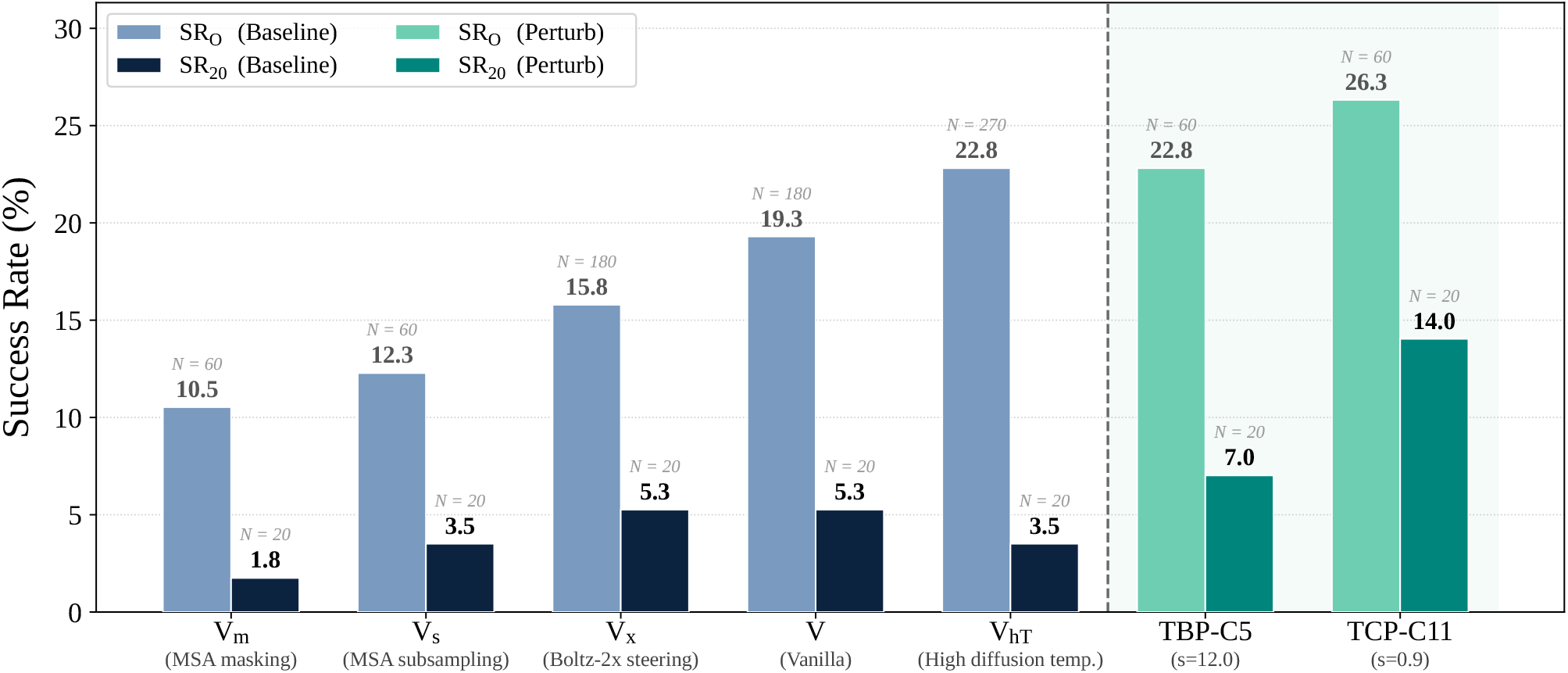
Baseline vs. perturbation success rates. Oracle (SR_*O*_) and top-20 (SR_20_) success rate (%) over 57 targets from Table 2, comparing baselines (navy 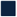 : *V, V*_*m*_, *V*_*s*_, *V*_*x*_, *V*_*hT*_) against perturbation methods (teal 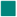 : TBP-C5, *s*=12.0; TCP-C11, *s*=0.9). *N* (above each bar) is the number of poses per target. See Table 2 for method definitions, metrics, and success criteria.

### 5.2. Confidence alone is not sufficient for pose selection

In the absence of ground truth, the confidence score predicted by the model serves as a proxy for prediction reliability. To assess how our perturbations affect the confidence score, we ranked samples for each PDB by confidence score, chose the top-20 confidence-scored samples per PDB, and then calculated the success rate for each PDB across the 20 highest confidence poses (Figure 5 and *SR*_20_ column in Table 2).

**Figure 5.**
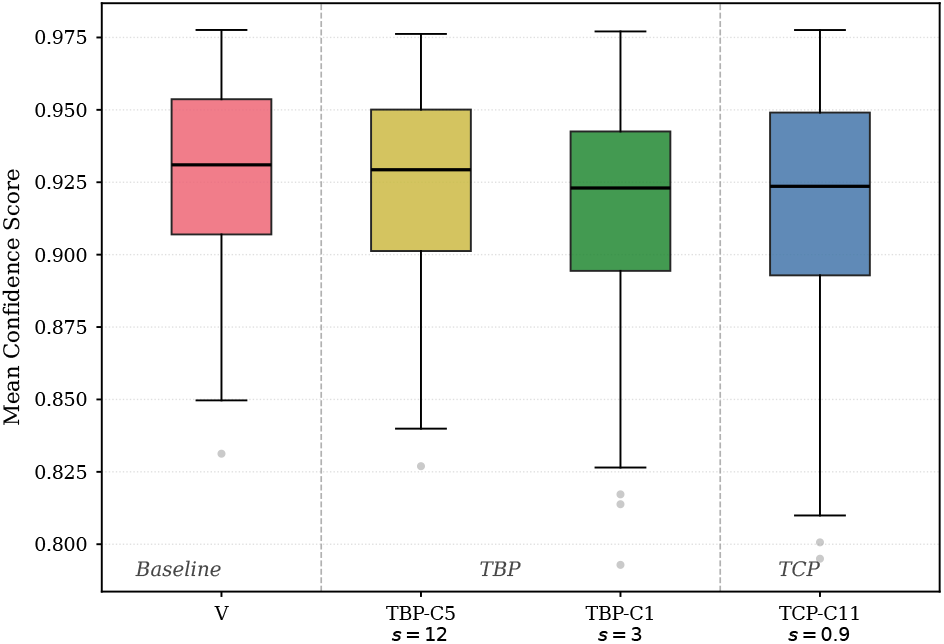
Per-target mean confidence score distributions on the RnP benchmark. For each of the 57 common PDB targets, the top20 poses (ranked by confidence score) are selected and their mean confidence score is computed. Box plots display the distribution of these per-target means across all methods. For the full distribution, refer to Figures S4 and S5.

The perturbation methods significantly outperformed all baseline methods. Notably, TCP-C11 more than doubled the success rate of the best baselines (5.26% to 14.04%). In Figure 5, we see that the overall mean confidence scores are similar across experiments and methods. This means that perturbation methods achieve higher success rates at comparable confidence levels, suggesting that confidence-based filtering is a more effective selection strategy for perturbation ensembles than for vanilla ensembles (*SR*_20_ column in Table 2). However, the decoupling of confidence from structural accuracy highlights that improvement in model confidence is needed for downstream analysis.

## 6. Discussion

In this work, we introduce Boltz-Perturb, a training-free perturbation approach that modifies Boltz-2’s denoising module to improve sampling diversity and structural accuracy. Our coordinate injection experiments reframed the core challenge from a model accuracy problem to a sampling deficiency problem, amenable to inference-time solutions. Among the approaches tested, TBP was slightly more effective in diagnostic systems while TCP outperformed on the RnP benchmark. On the RnP benchmark specifically, the integration of TCP, TBP, and vanilla Boltz yielded up to a two-fold improvement in success rate relative to vanilla Boltz alone. TCP achieves 15% higher oracle success compared to the best vanilla Boltz variant, V_hT_, with 78% less sampling, which translates to approximately 78% less compute, and TBP achieves the same oracle success rate as V_hT_ with 78% less sampling (Figure 4). To our knowledge, this is the first perturbation analysis of an AF3-style co-folding architecture for small-molecule binding mode diversity. However, for targets in the lowest similarity regime, inference-time perturbation alone may not overcome the model’s limitation, suggesting that targeted fine-tuning or additional training data may be necessary.

We observed that optimal perturbation regions depended on error magnitude. High-error targets benefited most from perturbing protein–ligand cross-attention, while moderately accurate targets improved with additional ligand self-attention perturbation. Moreover, model confidence scores did not reliably reflect coordinate-level improvements, suggesting that complementary post-hoc methods such as physics-based re-scoring may be needed for pose selection. We further discuss limitations of confidence scores and the benefit of investigating multiple predictions with a perturbation strategy when ground truth structures are not available (see Supplemental Information, Section D.7). TADS-Auto showed promising results for TBP configurations, suggesting it may reduce reliance on manual hyperparameter tuning. While not always optimal, it provides a good starting point when the appropriate perturbation schedule is unknown.

Since Token Bias Perturbation (TBP) operates on large internal tensors and requires tracking diffusion state throughout the sampling process, it incurs additional memory overhead, although the computational cost of the perturbation itself is minimal. As a result, a small number of targets could not be evaluated on NVIDIA Tesla V100 32GB GPUs due to memory constraints and required GPUs with more memory such as NVIDIA A100’s or H100’s. Nevertheless, perturbation methods achieved higher success rates using only one-third of the sampling budget compared to the baseline.

We acknowledge limitations that we leave as future work, including additional exploration of hyperparameters to enable more complex noising schedules and possible application of non-gaussian noise. While important, quantitative binding free energy prediction and improvement of confidence metrics are also beyond the scope of our current work.

We have shown that correct binding modes are often encoded within co-folding models but are not recovered at prediction time because of limited sampling diversity. Targeted inference-time perturbations of the conditioning signals that influence the structure-generation module provide an effective strategy for improving binding-mode discovery without retraining, increasing the utility of structure-generation models for structure-based drug discovery. We look forward to further improving the Boltz-Perturb framework as well as future reports of impact on downstream drug discovery.

## 7. Acknowledgements

We thank our colleagues for helpful discussions. This research was fully funded by Merck Sharp & Dohme LLC, a subsidiary of Merck & Co., Inc., Rahway, NJ, USA.

## 8. Data and Code Availability

The datasets used in this study are all publicly available as detailed in the *Experiments* section. Runs N’ Poses (RnP) is available from Zenodo (https://zenodo.org/records/18366081) and the five diagnostic structures (9JF4, 9M4Q, 9PY4, 9RAY, 9Z1L) are available from RCSB (https://www.rcsb.org/). The perturbation algorithms and functions introduced in this work (TCP, TBP, TADS-Auto) are fully specified in Algorithms 1, 2, and S1. Boltz-2 is available on GitHub (https://github.com/jwohlwend/boltz). Our Boltz-Perturb implementation integrated into Boltz-2 is available at https://github.com/MSDLLCpapers/boltz-perturb.

## A. Background

### A.1. AF3-Style Co-Folding Model Architectures

AlphaFold 3 established a new standard for sequence-to-structure prediction across biomolecule modalities. Several open-source implementations have followed, including the Boltz family, Chai, OpenFold-preview, and Protenix (Abramson et al., 2024; Wohlwend et al., 2024; Passaro et al., 2025; Chai Discovery Team et al., 2024; The OpenFold3 Team, 2025; Protenix Team et al., 2026). Despite differences in training data, model scale, and implementation details, these models share a common architectural pattern, consisting of trunk modules for sequence information followed by a denoising module for per-atom structure generation. Single and pairwise representations computed from the trunk are used to condition a diffusion process.

#### Trunk

The trunk distills evolutionary and pairwise sequence information primarily from protein multiple sequence alignments (MSA), and in some variants, pre-trained protein language models (pLMs), into rich representations. In Boltz-2, the trunk consists of an MSA module and 64 PairFormer layers with triangular attention operations following the AF3 architecture (Passaro et al., 2025). The PairFormer iteratively refines both single and pair representations through triangle multiplication, triangle attention, and single attention with pair bias.

#### Denoising module

The diffusion-based denoising module iteratively refines atomic coordinates through a reverse diffusion process, conditioned on trunk representations and optionally user-provided constraints. It consists of an atom encoder, token transformer, and atom decoder. The token transformer contains 24 layers of 16-head multi-head attention. Trunk-derived representations are calculated once before processed through conditioning projection and remain fixed throughout the diffusion process, bridging sequence information from trunk to the structure module (Abramson et al., 2024; Wohlwend et al., 2024; Passaro et al., 2025; Chai Discovery Team et al., 2024; The OpenFold3 Team, 2025; Protenix Team et al., 2026). Conditioning enters through two pathways:

- **Single representation** *s*: enters via adaptive layer normalization (AdaLN)—providing learned scale and additive shift—and sigmoid gating, which multiplicatively modulates token activations. Both mechanisms act before query, key, and value projections, meaning *s* indirectly influences downstream attention computations. The diffusion timestep embedding is incorporated into *s* via addition.
- **Pairwise representation** *z*: is normalized and linearly projected into a per-head additive bias *B* on the attention logits, producing a query and key (*QK*^⊤^ term) independent shift that modulates token-token interactions regardless of query and key magnitudes. This bias encodes pairwise evolutionary and structural priors and remains fixed throughout all diffusion steps.

In the multi-head attention mechanism, the attention map for the *l*-th layer and *h*-th head is denoted by *A*_*l,h*_ (*Q, K*), computed by query and key projections (*Q*_*l,h*_, *K*_*l,h*_), where 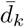 is the per-head dimensionality. Its output *O*_*l,h*_ is obtained by multiplying the attention map with the value projection *V*_*l,h*_. AF3-style co-folding models contain an explicit additive bias *B* in the attention computation that is absent from conventional image diffusion models.

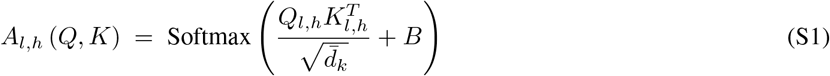

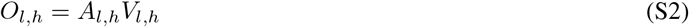

We evaluate on Boltz models, but the perturbation targets exist in all AF3-style architectures. Our method extends diversity improvement methods developed in other domains to the biomolecular setting.

#### Confidence module

A confidence module estimates prediction reliability, trained on mini-rollouts from the denoising module. In Boltz-2, this module uses 8 PairFormer layers on the final pair-token representations and encodings of the predicted coordinates, producing pairwise confidence estimates (PAE, PDE) with separate prediction heads for intra-chain and inter-chain pairs, from which ipTM is derived (Passaro et al., 2025). This is a lighter architecture than Boltz-1’s confidence module, which used a full 48-layer trunk.

### A.2. Conditioning signal perturbation

Our perturbation framework builds on CADS (Sadat et al., 2023), which demonstrated that adding scheduled Gaussian noise to conditioning signals during inference increases output diversity without retraining. CADS computes a time-annealed perturbed condition (*ŷ*) inspired by a forward diffusion process:

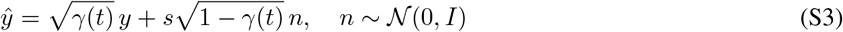

By Bayes’ rule, the conditional score decomposes as:

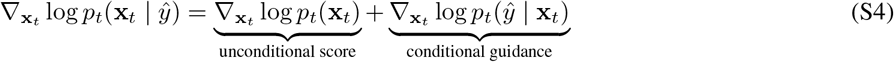

where **x**_*t*_ is the noisy data sample at diffusion step *t*. When *γ*(*t*) = 0, the perturbed condition is pure noise independent of **x**_*t*_, so the guidance vanishes and the model follows only the unconditional score.

As shown in (Sadat et al., 2023), for a differentiable denoiser *D*_*θ*_, this perturbation is equivalent to injecting non-isotropic Langevin noise into the sampling dynamics:

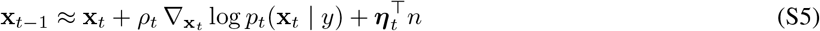

where 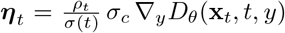. CADS showed that this strategy changes sampling dynamics more effectively than merely weakening guidance strength (Dynamic CFG) (Sadat et al., 2023). We adopt this principle for our perturbation targets with two key extensions. First, application to the pair and single representations of an AF3-style architecture.TCP is the direct analogue of CADS in our setting. The single representation *s* serves the same functional role as the conditioning signal *y* in CADS. Note that CADS derives a ridge regularization interpretation under a linear condition-to-data mapping assumption. In our setting, the conditioning pathway is non-linear, so this specific result does not formally hold. However, the qualitative insight—that conditioning noise smooths the score landscape and prevents mode dominance—is supported by our empirical results. Second, an adaptive scheduling strategy (TADS-Auto, Algorithm 1) that removes the need for manually specified injection windows by monitoring the model’s own stochastic noise at each step. For example in TCP, noise is added to the single-representation conditioning *s*_*t*_ on ligand tokens with a time-adaptive schedule (TADS, Eq. 3 or TADS-Auto). Our moment-matching rescaling step is analogous to the CADS rescaling operation that normalizes the perturbed condition back toward its original mean and standard deviation.

### A.3. Connection Between Perturbation Targets and Attention Mechanism

The single and pair representations enter the DiffusionTransformer through architecturally distinct pathways (Abramson et al., 2024). The single representation **s**_*i*_—which also carries the diffusion timestep embedding—conditions each block at four points: via AdaLN (multiplicative gating and additive shifting of token activations) before both the attention and transition sub-blocks, and via sigmoid output gating after each. Since queries, keys, and values are all derived from the AdaLN-modulated activations, perturbing **s**_*i*_ affects both token content and effective attention routing through modified queries and keys. The pair representation **z**_*ij*_ is projected into an additive bias on the pre-softmax attention logits, once per block, shifting the attention distribution independently of query and key content.

Both perturbation targets act exclusively within the DiffusionTransformer (24 blocks), leaving the AtomAttentionEncoder and Decoder unmodified. This restricts stochastic exploration to token-level structural decisions. While TBP and TCP enter through distinct pathways, their downstream effects are not fully separable: perturbations propagate across depth, and TCP’s modulation of queries and keys indirectly affects routing. We therefore treat the choice between TCP, TBP, and their combination as an empirical question.

#### Perturbation guidance in diffusion models

Beyond conditioning perturbation, several methods target the attention mechanism directly. SEG applies Gaussian blurring to attention logits, provably reducing the curvature of the energy landscape and producing smoother score estimates (Hong, 2024). PAG replaces attention maps with identity matrices to generate negative examples, while HeadHunter extends this to per-head perturbation for fine-grained control (Ahn et al., 2025b;a). Our approach differs in that we perturb the conditioning signals that feed into attention rather than the attention computation itself, allowing the model’s learned attention mechanism to translate conditioning noise into structurally coherent diversity. When we evaluated perturbations on head-level attention weight perturbation in token transformer, we did not get meaningful improvements on our diagnostic set.

### A.4. Other Diversity Strategies

There are several other existing methods that can potentially increase the diversity: temperature and seed variation, MSA subsampling, fine-tuning, and steering potentials. (1) Model Level: Different temperature and seed variation provides limited diversity. (2) Trunk Level: MSA subsampling modifies the evolutionary signal input to the trunk. (3) Training: Fine-tuning on known binding modes modifies the model parameters to bias predictions toward specific conformations of interest but requires target-specific retraining and known reference structures. (4) Diffusion Level: Steering potential from Boltz-1, Boltz-2 applies physics-based potentials gradient signal to the predicted denoising coordinate during reverse diffusion.

The Boltz-steering method provides inference-time steering that biases reverse diffusion trajectories. Following the authors’ notation, *τ* (*x*_*t*_|*f*_*θ*_(*x*_*t*+1_, *t* + 1), *x*_*t*+1_, *t*) is a sampling algorithm used by Boltz at timestep t, and corresponding transition distribution is *p*_*θ*_(*x*_*t*_|*x*_*t*+1_) = *τ* (*x*_*t*_|*f*_*θ*_(*x*_*t*+1_, *t* + 1), *x*_*t*+1_, *t*). Boltz-steering is applied in the form of universal guidance by 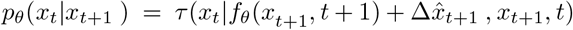 where 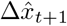 is computed by gradient descent with respect to added physics-based energy differences.

In contrast, our method does not optimize toward a specific objective function. Instead, it injects stochastic noise into the conditioning representations—pair bias and single-token modulation—within the DiffusionTransformer, encouraging exploration of alternative denoising trajectories without requiring target-state supervision. This places our approach closer to stochastic perturbation guidance than to directed optimization: rather than steering the model toward a predefined state, we broaden the distribution of sampled conformations by perturbing the conditioning signals.

## B. Methods

### B.1. Algorithm

This section describes the algorithm.

#### Algorithm S1

Boltz-Perturb Inference Pipeline

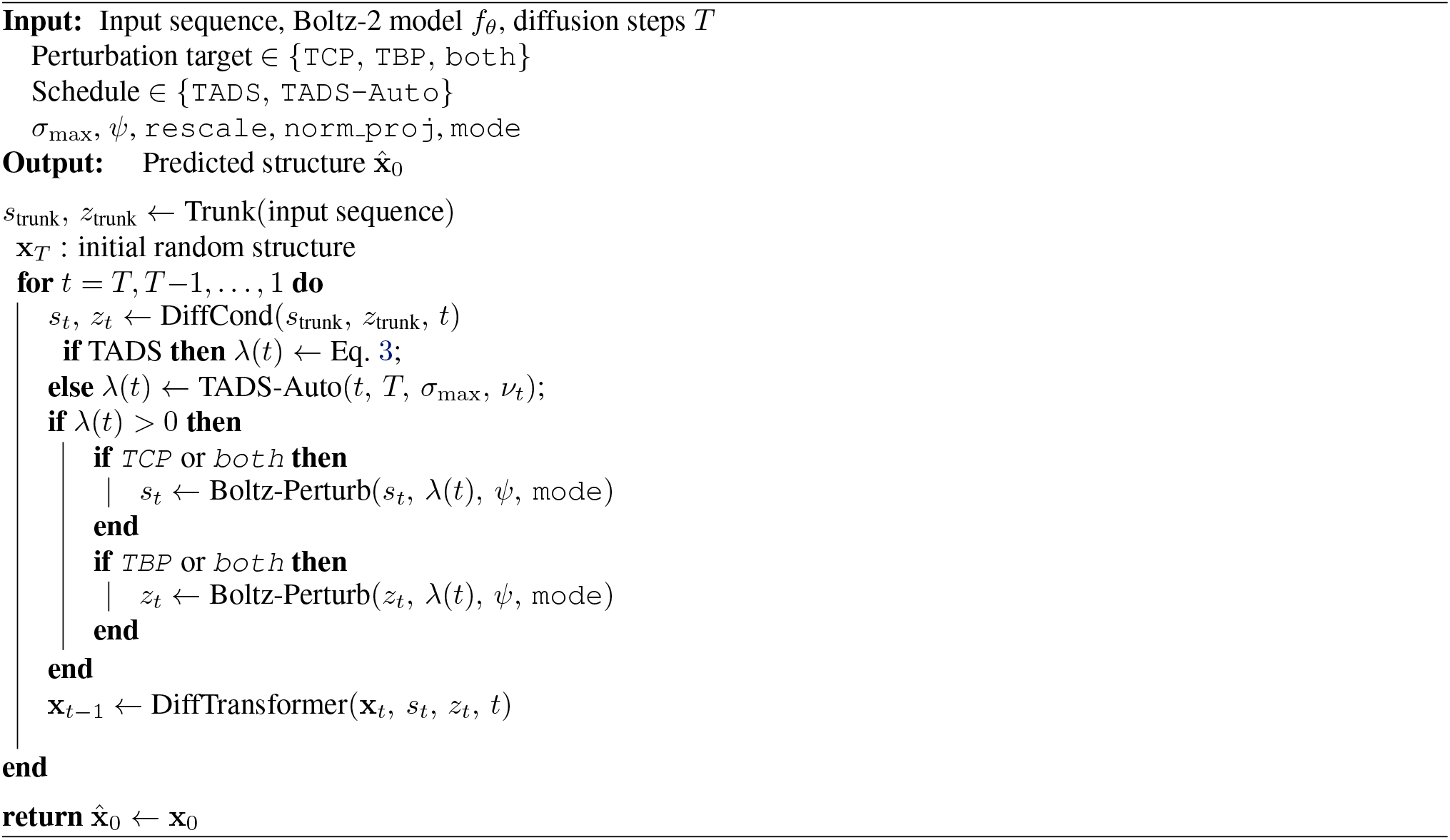

The single token representations are passed as *s*_trunk_ and *s*_inputs_. *s*_inputs_ is produced in embedder directly from the input feature dictionary, encoding per-token chemical identity (residue type, molecule type, bond connectivity) without any cross-token reasoning, whereas at the trunk-level *s*_trunk_ is the output of the Pairformer. At each diffusion step *t*, `SingleConditioning` combines the static trunk representation *s*_trunk_, the static input features *s*_inputs_, and the current timestep embedding *γ*(*t*) into a single per-token conditioning tensor *s*_*t*_; `SingleConditioning` is only dependent on the time *t*. Now, because *s*_trunk_ and *s*_inputs_ are replicated identically across all *M* samples before this call, all *M* copies of *s*_*t*_ are identical at every step.

Both *B* and *s*_*t*_ are derived from the trunk outputs and shared identically across all *M* denoising samples. *B* is fully static. It is fixed for all diffusion steps and all samples. *s*_*t*_ varies across steps but remains sample-identical: at any given step *t* all *m* samples receive the same *s*_*t*_. We hypothesized that this sample-identity or static nature acts as a diversity bottleneck. All *m* denoising trajectories are conditioned on the same relational and contextual prior. TBP and TCP break this by making *B*^(*m*)^ and 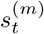 sample-specific for each *m* ∈ *{*1, …, *M*}.

### B.2. Additional dataset

#### Diagnostic set

The 5 complexes were selected from PDB depositions after the RnP benchmark curation date and Boltz-2 training cutoff, with the criteria: (1) single protein chain and (2) single non-covalent ligand, PDB IDs and ligand identifiers are listed in Table 1.

#### Curated Runs N’ Poses benchmark set

We assessed Boltz-Perturb using the Runs N’ Poses (RnP) benchmark. Starting from the full RnP benchmark of RnP, we selected test systems containing a single protein chain and a single ligand molecule released after the Boltz-2 training cutoff date. We then retained only PDBs in which the RnP reported the best Boltz-2 predicted ligand RMSD exceeded 2Å, ensuring that our evaluation focused on cases where Boltz-2 did not already produce an accurate ligand pose. We observed that higher perturbation noise can shift predicted ligand positions outside the conventional binding pocket, implying the potential to discover alternative binding sites.

### B.3. Baseline Details

#### Vanilla Boltz-2 (V)

Baseline conditions were generated using Boltz-2 with default parameters: 3 random seeds, 2 sampling temperatures, and 30 diffusion samples per seed-temperature combination, yielding 180 poses per target.

#### Steering potentials (Vx)

Boltz with steering potential (use potentials) flag

#### Elevated temperature (V high-T)

Diffusion temperature *T* ∈ *{*1.2, 1.3, 1.4*}*

#### MSA masking (V mask)

60 samples. Random masking of MSA columns at rate 0.1.

#### MSA subsampling (V sub)

60 samples. MSA depth reduced to 4086 rows.

### B.4. True Coordinates Insertion Experiment

We prepared a structure coordinate file from the PDB as a NumPy array. During the denoising process, if the noise variance was lower than a set threshold, we replaced the coordinates with the prepared PDB coordinates once. Denoising then continued following the default noise schedules. This was similar to resampling, in that the true coordinates were injected, then noised and denoised according to the pre-scheduled default noise schedules. During the true coordinate experiments, some runs successfully recovered the correct binding mode even when noise was added. However, some did not recover as well. For example, we observed flipped conformations. This experiment led us to conclude that the focus needed to be on reaching the correct conformational space and on how to get there efficiently. The use of confidence scores was not meaningful, as the scores were not consistent across runs.

## C. Metric Definitions

### C.1. RMSD (Root Mean Square Deviation) Formula

We first align the predictions to the ground truth structure.

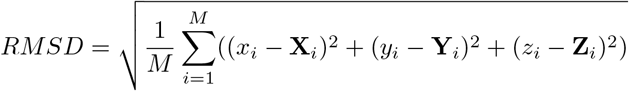

where *M* is the number of atoms, (*x*_*i*_, *y*_*i*_, *z*_*i*_) are the (*x, y, z*) coordinates of the *i*-th atom in the predicted structure respectively, and (**X**_*i*_, **Y**_*i*_, **Z**_*i*_) are the coordinates of the *i*-th atom in the reference (ground truth) structure.

### C.2. RMSF (Root Mean Square Fluctuation) Formula

#### No Protein Context RMSF

For each ligand atom *i*, we compute the Root Mean Square Fluctuation (RMSF) across *N* samples as follows:

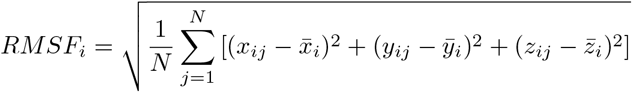

where (*x*_*ij*_, *y*_*ij*_, *z*_*ij*_) are the coordinates of atom *i* in sample *j*, and 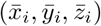 are the mean coordinates of atom *i* across all samples *N*. The mean coordinates are computed using RDKit aligned to the first frame.

#### Protein Context RMSF

For each sample, we use PyMOL to align the protein in the complex to the ground truth structure, producing aligned ligand coordinates. Then, for each **aligned** ligand atom *i*, we compute the Root Mean Square Fluctuation (RMSF) across *N* samples as follows:

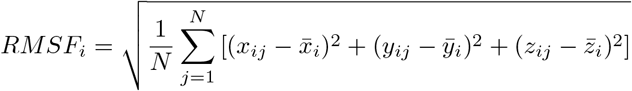

where (*x*_*ij*_, *y*_*ij*_, *z*_*ij*_) are the coordinates of atom *i* in sample *j*, and 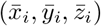 are the mean coordinates of atom *i* across all samples *N*. The mean coordinates are computed using the protein-aligned ligand coordinates, which are obtained by aligning the protein in each sample to the ground truth structure, and then applying the same transformation to the ligand coordinates.

#### Mean RMSF

After individual RMSF values are computed for each ligand atom, we calculate the mean RMSF across all ligand atoms to obtain an overall measure of fluctuation:

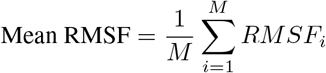

where *M* is the total number of ligand atoms. This mean RMSF provides a single number.

#### Success rate (SR)

Fraction of unique test systems, where the minimum ligand RMSD from openstructure is lower than 2Å

#### Structural validity

All generated structures were filtered for validity using PoseBusters (v0.6.5) (Buttenschoen et al., 2024) and OpenStructure (v2.7.0). Invalid predictions were excluded from downstream analysis. PoseBusters checks: bond lengths, bond angles, internal clashes, protein–ligand clashes, volume overlap. A pose is considered valid if all checks are satisfied.

#### Confidence scores

Boltz-2 reported confidence score as model self-assessment.

## D. Additional Results and Discussion

### D.1. Hyperparameter Grid

Table S1 lists all perturbation conditions evaluated.

**Table S1.**
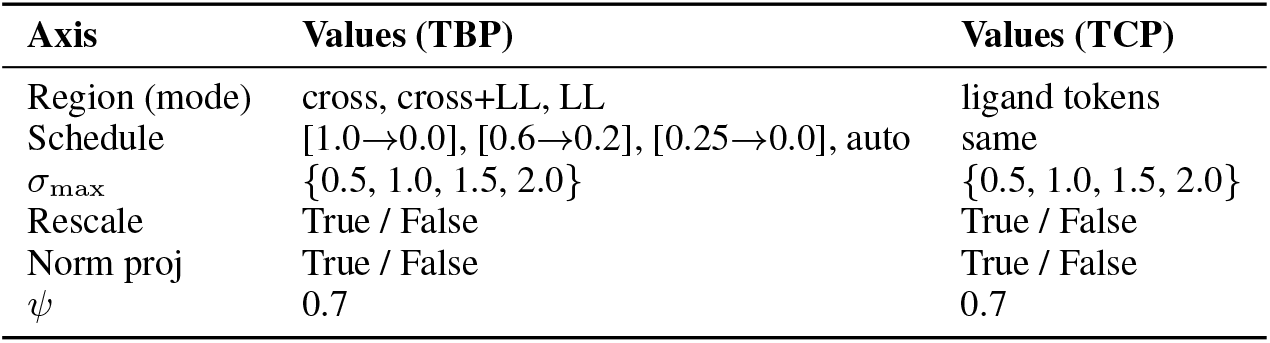
Full hyperparameter grid for TBP and TCP experiments.

#### region-specific masking

We applied region-specific masking schemes to protein and ligand tokens to disentangle their individual and joint contributions to binding mode prediction. Specifically, we evaluated four masking configurations: no masking (PP+PL+LP+LL), ligand self-attention only (LL), protein–ligand cross-attention only (PL+LP), and all regions except protein self-attention (PL+LP+LL).

#### Noise schedules

We evaluated four noise scheduling strategies: three fixed injection windows—full trajectory (*t* ∈ [0.0, 1.0]), mid-range (*t* ∈ [0.2, 0.6]), and tail-end (*t* ∈ [0.0, 0.25])—as well as the adaptive TADS-Auto schedule. The tail-end window was selected based on the empirical observation that, in the 200-step diffusion process, the stochastic noise variance effectively vanishes around step 154 (corresponding to *t* ≈ 0.25).

#### Regularization ablation

To assess the impact of regularization, we independently ablated moment-matching rescaling (R) and norm-preserving projection (N) under a fixed full-trajectory injection schedule.

#### Sampling

For each perturbation condition and noise setting, we generated 60 samples using the same random seeds and temperature combinations as vanilla Boltz-2, but with 10 samples per condition instead of 30.

### D.2. Vanilla and True Coordinate Injection Outputs

**Figure S1.**
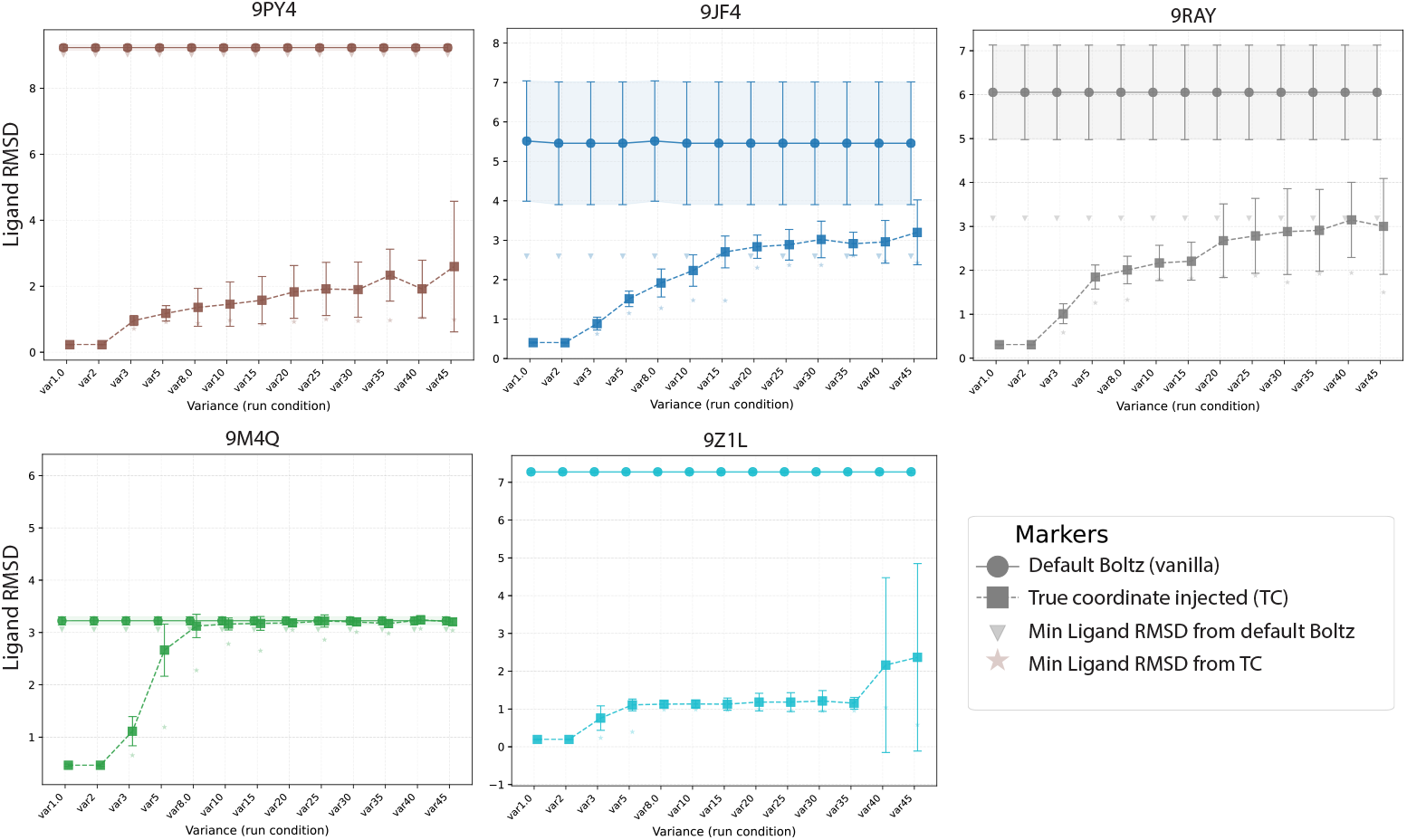
Diagnostic five set true coordinate injection outputs.

**Figure S2.**
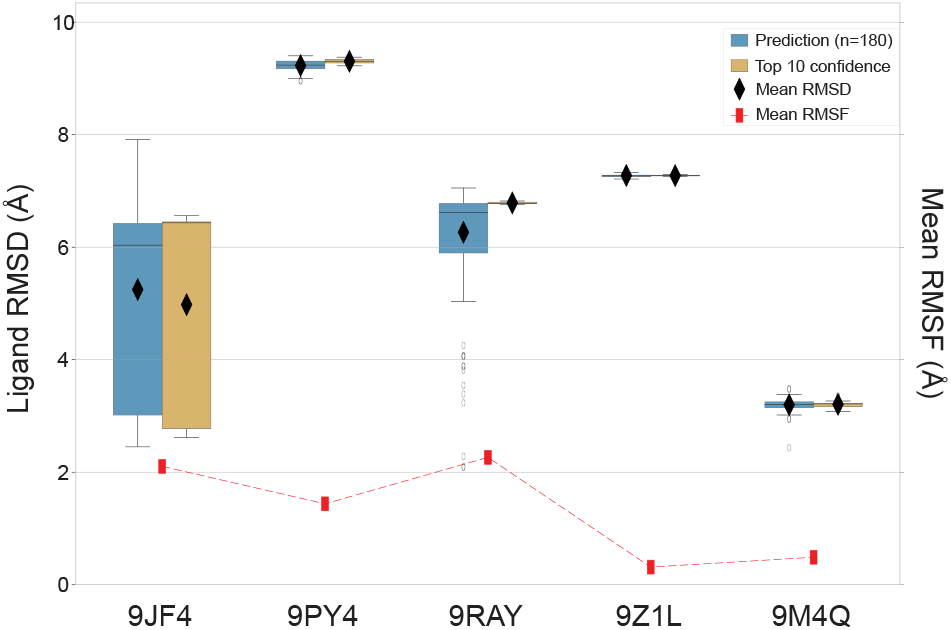
Diagnostic five set vanilla performance.

### D.3. More Results of Perturbation in Different Conditions

**Table S2.** Minimum ligand RMSD (Å) per method across all 5 PDB targets (best over all noise scales). TBP/TCP values shown as *RMSD (ns)*. Bold = column minimum. `---` = no valid pose.

| Method | 9JF4 | 9M4Q | 9PY4 | 9RAY | 9Z1L |
| --- | --- | --- | --- | --- | --- |
| V | 2.45 | 2.43 | 8.96 | 2.09 | 7.21 |
| V<sub>x</sub> | 2.25 | 3.01 | 9.01 | 3.82 | 7.21 |
| V<sub>highT</sub> | 2.25 | 3.01 | 9.01 | 3.82 | 7.21 |
| V<sub>mask</sub> | 2.87 | 3.02 | 8.98 | 1.99 | 7.21 |
| V<sub>sub</sub> | 2.34 | 3.06 | 9.02 | 4.00 | 7.21 |
| TBP_C1 | 2.53 (ns=1.0) | **1.90 (ns=3.0)** | 8.69 (ns=5.0) | 2.04 (ns=0.1) | 6.87 (ns=5.0) |
| TBP_C2 | 2.37 (ns=1.0) | 2.90 (ns=20.0) | **1.67 (ns=3.0)** | 3.24 (ns=17.0) | 1.15 (ns=20.0) |
| TBP_C3 | 2.11 (ns=3.0) | 1.92 (ns=3.0) | 8.79 (ns=4.0) | 2.24 (ns=0.1) | 6.88 (ns=5.0) |
| TBP_C4 | 2.50 (ns=1.0) | 2.37 (ns=9.0) | 2.03 (ns=11.0) | **1.06 (ns=25.0)** | 1.07 (ns=12.0) |
| TBP_C5 | 2.47 (ns=0.3) | 2.64 (ns=10.0) | 8.98 (ns=0.8) | 1.93 (ns=10.0) | 7.14 (ns=10.0) |
| TBP_C6 | 2.76 (ns=1.0) | 2.09 (ns=11.0) | 8.31 (ns=20.0) | 3.64 (ns=1.0) | 6.67 (ns=20.0) |
| TBP_C7 | 2.13 (ns=1.0) | 1.91 (ns=2.0) | 8.71 (ns=2.0) | 2.05 (ns=0.3) | **6.86 (ns=3.0)** |
| TBP_C8 | 2.28 (ns=2.0) | 1.99 (ns=3.0) | 8.43 (ns=3.0) | 3.20 (ns=1.0) | **6.86 (ns=3.0)** |
| TBP_C9 | 2.56 (ns=1.0) | 2.34 (ns=14.0) | 9.01 (ns=8.0) | 1.81 (ns=25.0) | 7.15 (ns=25.0) |
| TCP_C10 | 2.23 (ns=0.6) | 2.67 (ns=0.2) | 1.86 (ns=0.6) | 2.06 (ns=0.7) | **0.97 (ns=0.7)** |
| TCP_C11 | **1.76 (ns=1.0)** | 2.33 (ns=1.0) | 1.96 (ns=0.9) | 1.99 (ns=0.9) | 1.07 (ns=0.7) |
| TCP_C12 | 2.37 (ns=0.8) | 2.92 (ns=0.9) | 9.02 (ns=0.8) | 5.47 (ns=0.5) | 7.18 (ns=1.0) |

V: Vanilla Boltz; V<sub>mask</sub>: MSA mask; V<sub>sub</sub>: MSA subsample. TBP: Token Bias Perturbation; TCP: Token Conditioning Perturbation.
C1: PL+LP+LL, $t \in [0.0, 1.0]$ ; C2: PL+LP, $t \in [0.2, 0.6]$ ; C3: PP+PL+LP+LL, $t \in [0.0, 1.0]$ ; C4: PL+LP, $t_{\text{auto}}$ ; C5: PL+LP, $t \in [0.0, 1.0]$ ;
C6: PL+LP+LL, $t \in [0.2, 0.6]$ ; C7: PL+LP+LL, $t \in [0.0, 1.0]$ , $\psi=1.0$ ; C8: PL+LP+LL, $t_{\text{auto}}$ ; C9: PL+LP, $t \in [0.2, 0.6]$ , physical guidance; C10: $t \in [0.0, 1.0]$ (TCP variant) C11:
L $t \in [0.2, 0.6]$ (TCP variant). C12: L $t_{\text{auto}}$ (TCP variant).

#### D.3.1. Diagnostic Five

We also report the validity trend for the experiments.

From Figure S3, the experiments that include LL (ligand region), tend to drop in validity as the noise scale increases.

**Figure S3.**
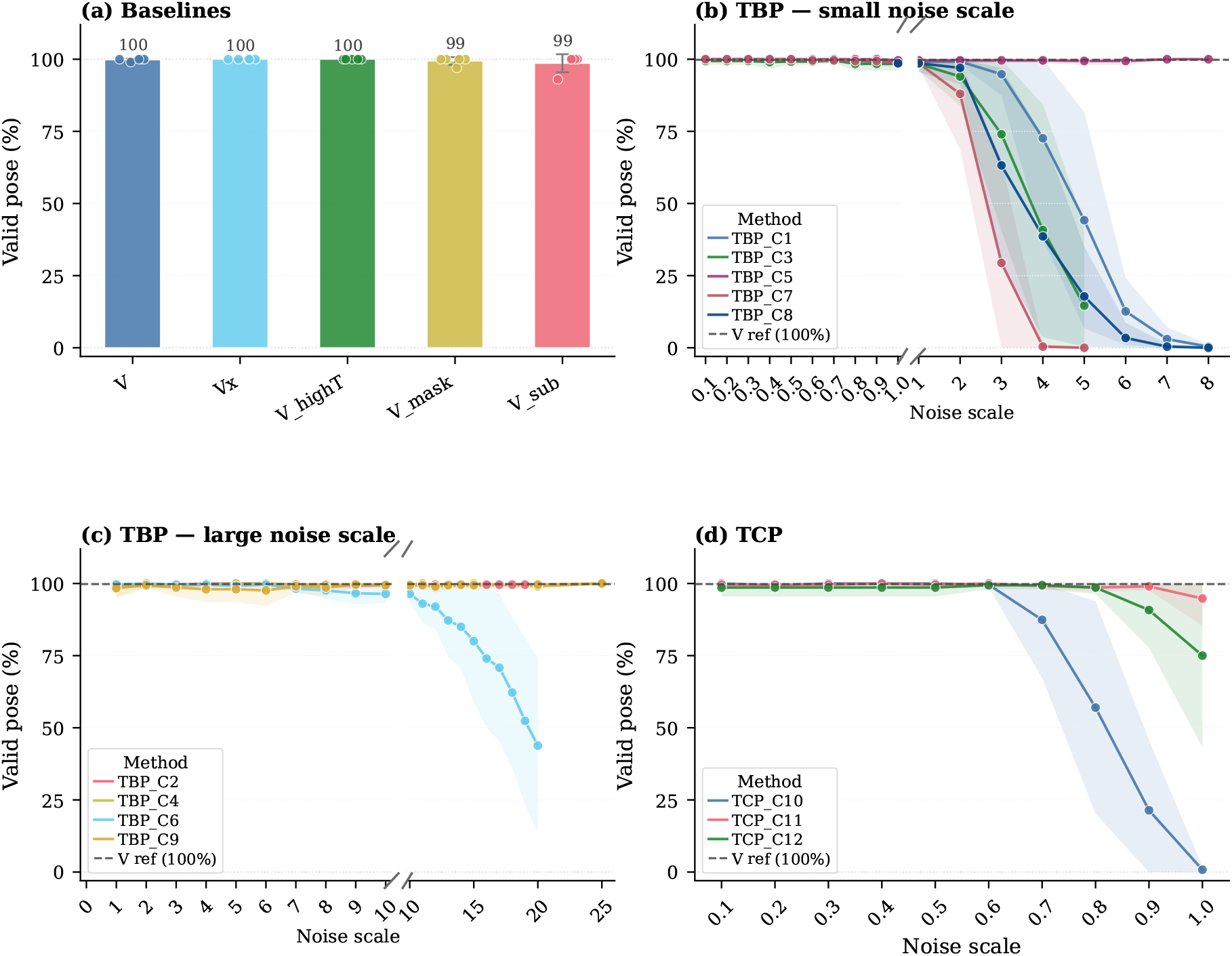
Valid pose percentage across perturbation experiments. For each noise scale, the percentage of valid structures (validity) is averaged across all five PDB targets. **(a)** Baselines: mean valid pose % per method averaged across five PDB targets (bars), with individual target values overlaid as dots and error bars denoting *±*1 std. **(b)–(d)** Mean validity % *±* 1 std (shaded) across five PDB targets as a function of noise scale *σ*: **(b)** TBP small-scale (*σ* ∈ [0.1, 8]), **(c)** TBP large-scale (*σ* ∈ [0, 25]), and **(d)** TCP (*σ* ∈ [0.1, 1.0]). The dashed line indicates the Vanilla (V) reference mean.

### D.4. Confidence Results of TBP

We report the confidence score distribution for two different datasets.

#### D.4.1. Diagnostic Five

**Figure S4.**
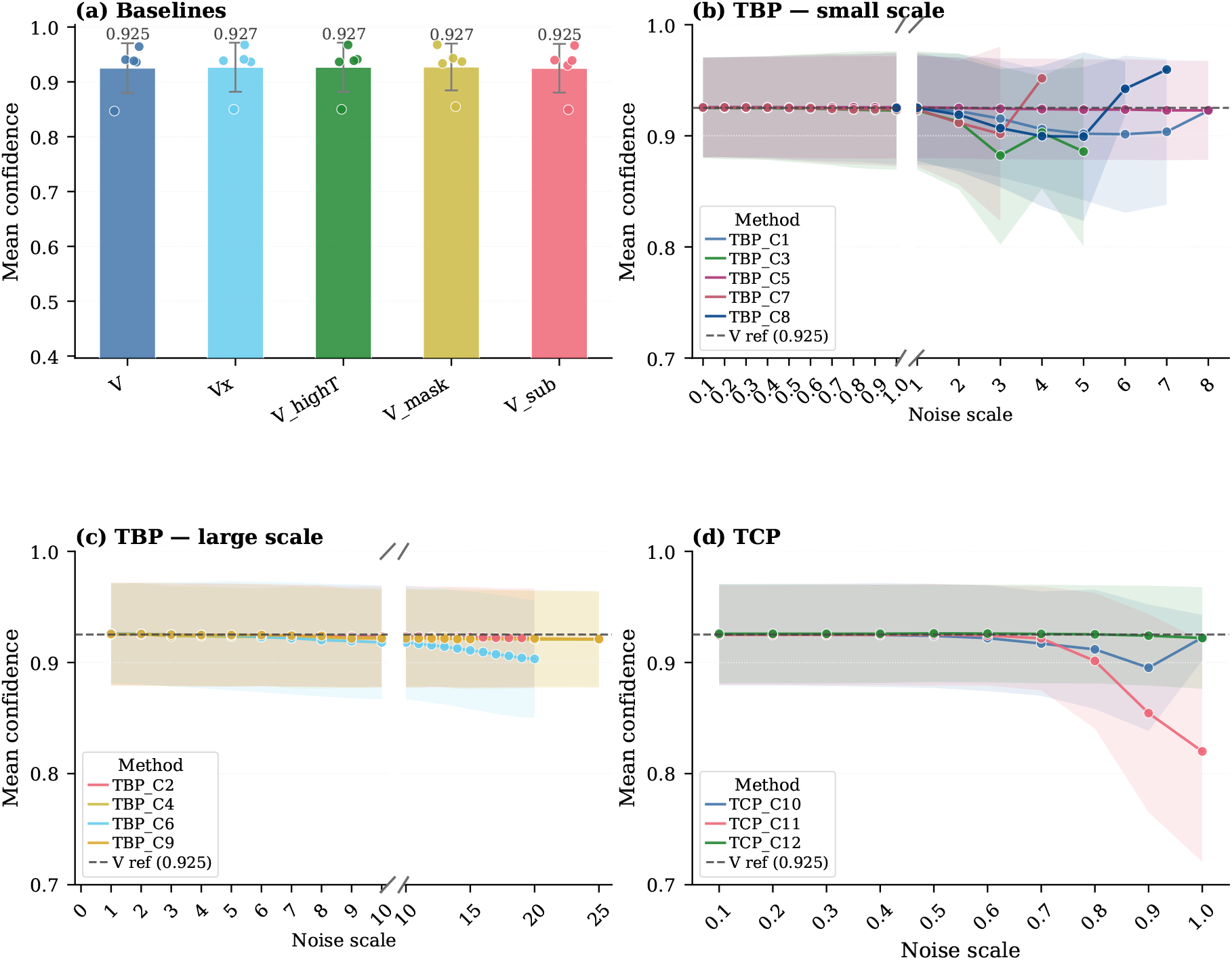
Mean confidence score as a function of noise scale *σ*_*max*_ across perturbation experiments (only the valid structures). **(a)** Baseline methods: bars show the mean confidence averaged across PDB targets; individual target values are overlaid as dots; error bars denote *±* 1 std across targets. **(b)–(d)** Each point represents the mean-of-means confidence score: for each target, confidence scores are first averaged across all poses at a given *σ*_*max*_ (level 1 mean); these per-target means are then averaged across all PDB targets (level 2 mean), with *±*1 std shaded. The dashed line marks the Vanilla (V) reference.

#### D.4.2. RnP Benchmark

**Figure S5.**
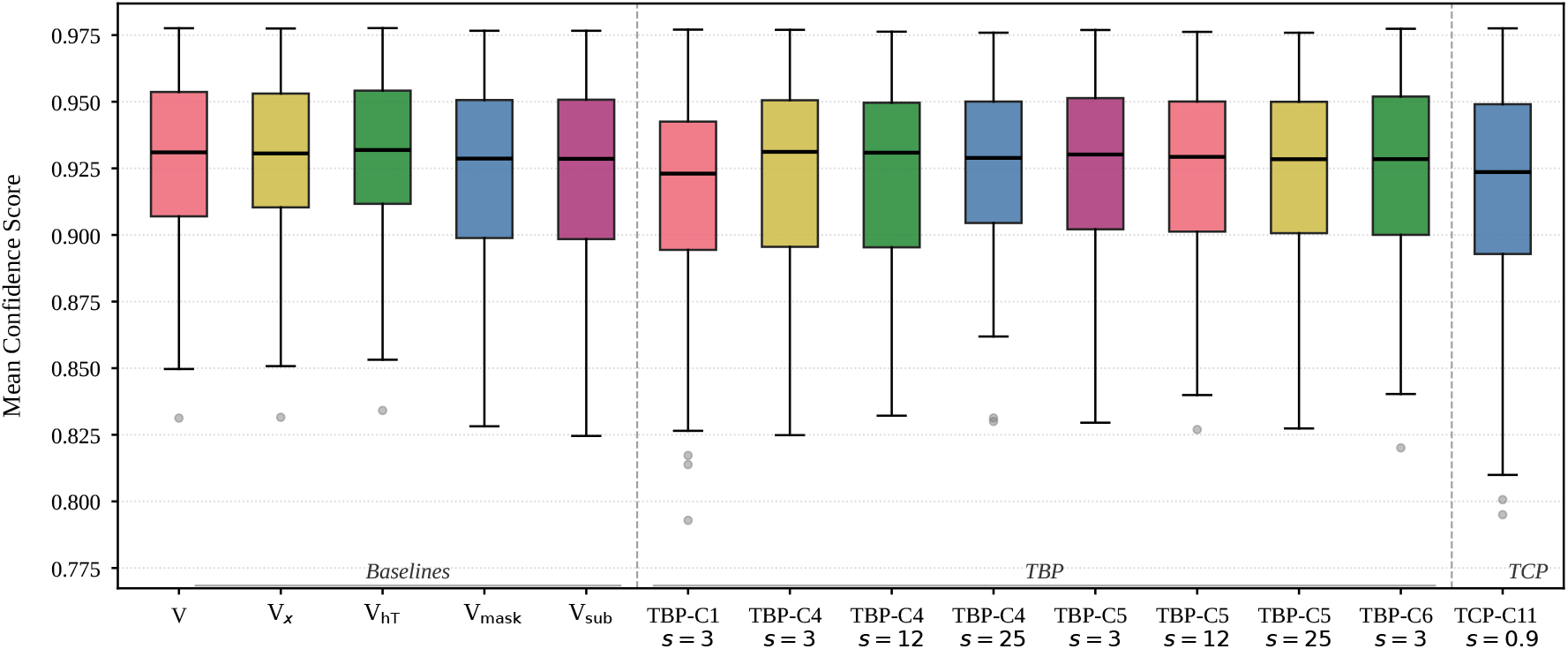
Per-target mean confidence score distributions on the RnP benchmark. For each of the 57 common PDB targets, the mean confidence score is computed over all generated poses. Box plots display the distribution of these per-target means across all methods.

### D.5. Common Failure Modes

Common pose prediction failures occur at multiple scales. At the local level, protein side chain and ligand torsion errors are common. Pocket failures range from small side chain rotamer errors—where a protein side chain is predicted in the wrong rotational conformation—to helix conformation changes and large-scale residue displacement.

#### D.5.1. GPU-Dependent Variability

Table S3 shows a concrete example of GPU-dependent variability for target 9PY4 under TBP. Despite identical software conditions, NVIDIA Tesla V100 32GB GPUs produced substantially different RMSD values (1.674 Å) compared to NVIDIA A100 80GB or NVIDIA H100 80GB (9.0 Å). The results within each GPU family remained highly consistent regardless of kernel configuration. The factors mentioned in section D.6 could have affected the results, but the exact cause remains unclear.

**Table S3.** RMSD (Å) for target 9PY4 across GPU types and kernel configurations under identical software conditions. All runs use the same CUDA version and random seed.

| Condition | H100 | H100 | A100 | A100 | V100 | V100 |
| --- | --- | --- | --- | --- | --- | --- |
|  | No Kernel | Kernel | No Kernel | Kernel | No Kernel | Kernel |
| PL+LP<sub>t$\in$[0.6, 0.2]</sub> | 9.167 | 9.164 | 9.009 | 9.007 | 1.674 | 1.674 |

### D.6. GPU Type Can Affect Performance

We observed that RMSD can vary across GPU types even when the conda environment, CUDA version, and precision settings are identical. In vanilla Boltz-2, this effect is negligible (Table S4). Under perturbation, however, the additional matrix multiplications, element-wise operations, and random noise injection introduce extra computation that different GPU architectures may execute differently, leading to small numerical differences. These differences may be due to, but are not limited to, bf16 rounding behavior across hardware generations, non-deterministic atomic operations, cuDNN runtime algorithm selection, whether tensor cores were enabled, and floating-point reduction ordering across streaming multiprocessors (PyTorch Contributors, 2024; NVIDIA Corporation, 2024a;b; Fasi et al., 2021), all of which are known sources of numerical non-determinism in deep learning inference and are not specific to our method (Schlögl et al., 2023; Yuan et al., 2025). Since each experiment was run on whichever GPU resources were available at the time of HPC cluster submission, the exact GPU type per run was not recorded. We report this transparently as a practical limitation of large-scale cluster-based experimentation and leave systematic characterization of GPU-dependent variability to future work.

#### D.6.1. Vanilla

**Table S4.** Minimum ligand RMSD (Å) per structure for each GPU. All runs use 32-bit precision.

| PDB | H100 | H100 | A100 | A100 | V100 | V100 |
| --- | --- | --- | --- | --- | --- | --- |
|  | No Kernel | Kernel | No Kernel | Kernel | No Kernel | Kernel |
| 9JF4 | 2.451 | 2.457 | 2.459 | 2.459 | 2.459 | 2.459 |
| 9M4Q | 2.433 | 3.013 | 3.017 | 3.017 | 3.017 | 3.017 |
| 9PY4 | 8.958 | 8.929 | 8.929 | 8.929 | 8.929 | 8.929 |
| 9RAY | 2.089 | 2.153 | 2.154 | 2.154 | 2.154 | 2.154 |
| 9Z1L | 7.212 | 7.192 | 7.192 | 7.192 | 7.207 | 7.207 |

The differences are negligible across different GPUs for vanilla.

### D.7. Practical Scenario: Pose Selection Without Ground Truth

Without access to ground-truth structures, researchers typically curate a diverse set of predicted structures above certain confidence threshold for downstream analysis (e.g., MD simulation, free energy calculations). Therefore, assessing performance such as ligand RMSD *<* 2Å based on top-*k* structures ranked by the confidence metrics reflects widely adopted evaluation protocol in the field. Here, we further illustrate the relationship between confidence and accuracy, which support that Boltz-Perturb can be useful in practice through improved sampling.

We used the results of the 57-target RnP benchmark from Table 2, but kept only targets with at least one correct pose (ligand RMSD *<*2Å) when combining all predictions. This yielded 23 out of 57 targets. We performed top-*k* confidence analysis between TCP-C11 (*s* = 0.9) and baseline variants–vanilla Boltz-2 (*V*), vanilla Boltz-2 with steering potential (*V*_*x*_)–from Table 2.

Per-target scatter plots compare the top-20 poses ranked by confidence metrics against their ligand RMSD, Figures S6 for confidence score and Figures S7 for ligand ipTM, respectively. The TCP-C11 spans a wider range in both confidence and ligand RMSD compared to baselines *V* and *V*_*x*_,consistent with increased sampling diversity. This supports our hypothesis that increased sampling diversity helps predict correct pose(e.g., 8ivr, 8sge from Figure S6 and 8sge, 8jot, 8pqa from Figure S7). TCP-C11 places poses below 2Å within its top-20 whereas the baselines have none or far fewer. For 8sge in particular, TCP-C11 yields many more correct poses (ligand RMSD *<* 2Å) across the confidence range. Ranking by ligand ipTM yields a similar result.

However, we caution that a high confidence metric value alone is not a reliable indicator of a correct pose. For example, the accurate TCP-C11 structures are not always the “single” most-confident prediction. This is why we report the top-30 ranked poses rather than trusting the top-1 and in this case TCP-C11 places more correct poses within that top-20.

In a real-world scenario, we would want multiple probable candidates within a given top-*k* rather than just a single one, as we do not know which will ultimately be correct. More options can increase confidence in the selection and provide more starting points for downstream analysis in drug discovery.

Figure S8 extends this analysis by varying *k*. TCP-C11 consistently achieves more predictions with RMSD *<* 2 Å at every *k* compared to all baselines, despite having fewer total samples as noted in Table 2. For the baseline to match TCP’s maximum count per target, which TCP-C11 achieves around *k* = 40, *V* (Boltz-2 vanilla) requires approximately 120 predictions (≈3*×*) and *V*_*x*_ requires approximately 90 (≈2.5*×*). Therefore, TCP-C11 does not merely diversify predictions—it increases the number of quality poses recoverable through confidence-based selection.

These results show that Boltz-Perturb’s diversity translates into practical utility in drug discovery: more correct poses are recoverable through confidence-ranked selection than those of baselines, without requiring additional predictions. A thorough analysis of the correlation between confidence metrics and pose quality (e.g., ligand RMSD) would be needed to clarify when confidence-based selection is reliable, and we leave this for future work.

**Figure S6.**
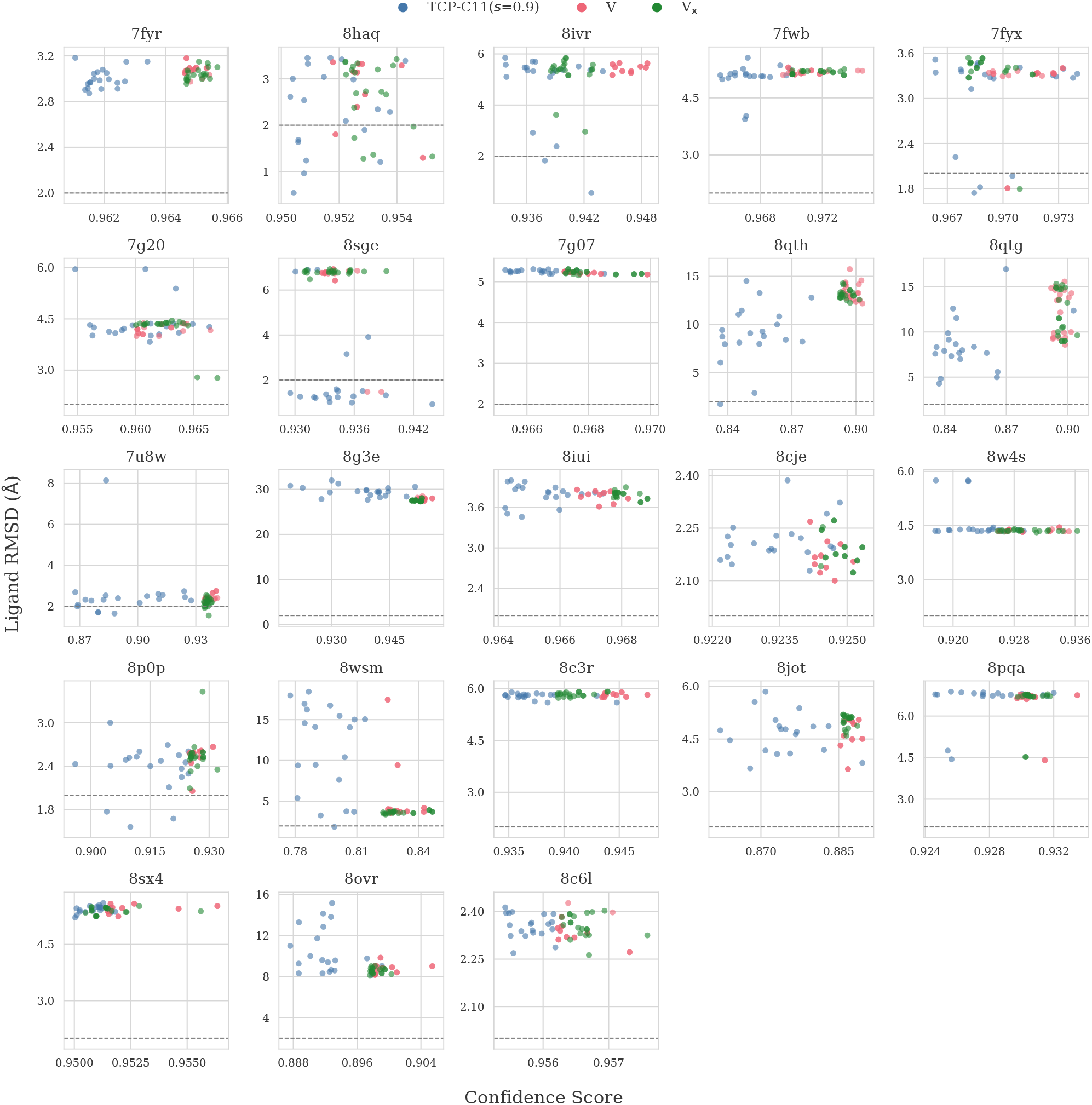
Whether high-confidence poses are also accurate, per target. Per-target confidence–accuracy (ligand RMSD) scatter. Each panel is one target comparing TCP-C11 against *V* (vanilla Boltz-2) and *V*_*x*_ (Boltz-2x steering potential); points show the top-20 poses *per method*, ranked by confidence score (`confidence_score`), on the x-axis against the ligand RMSD on the y-axis. Colors distinguish methods. The dashed grey line is the 2 Å threshold. Only targets with at least one pose below 2 Å ligand RMSD (oracle success) are included, giving *n* = 23 out of 57 targets. Axes are autoscaled. All 57 targets shown in Figure S9.

**Figure S7.**
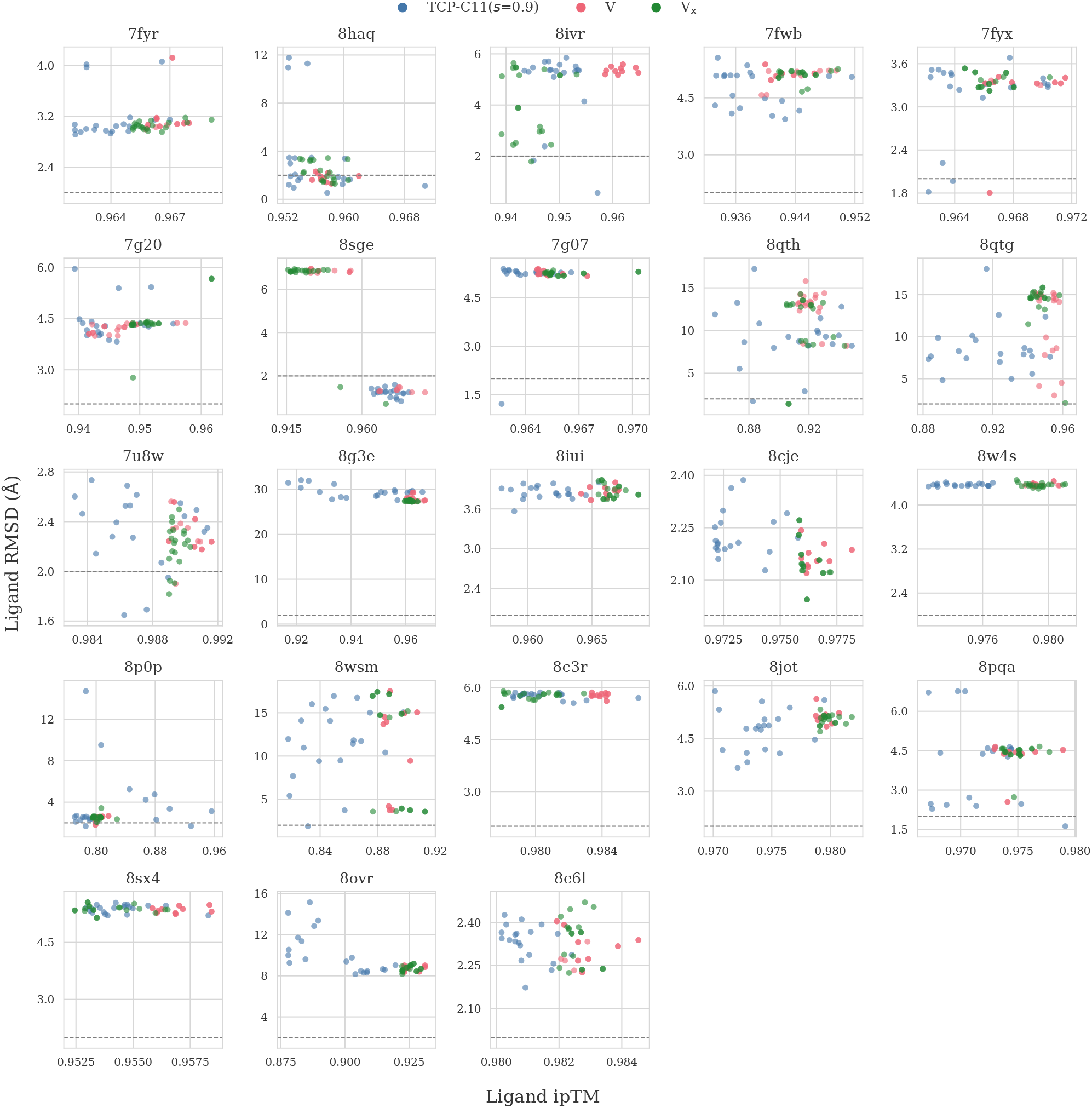
Whether high-ipTM poses are also accurate, per target. Per-target ipTM–accuracy (ligand RMSD) scatter. Each panel is one target comparing TCP-C11 against *V* (vanilla Boltz-2) and *V*_*x*_ (Boltz-2x steering potential); points show the top-20 poses *per method*, ranked by ligand ipTM (`ligand_iptm`), on the x-axis against the ligand RMSD on the y-axis. Colors distinguish methods. The dashed grey line is the 2 Å threshold. Only targets with at least one pose below 2 Å ligand RMSD (oracle success) are included, giving *n* = 23 out of 57 targets.

**Figure S8.**
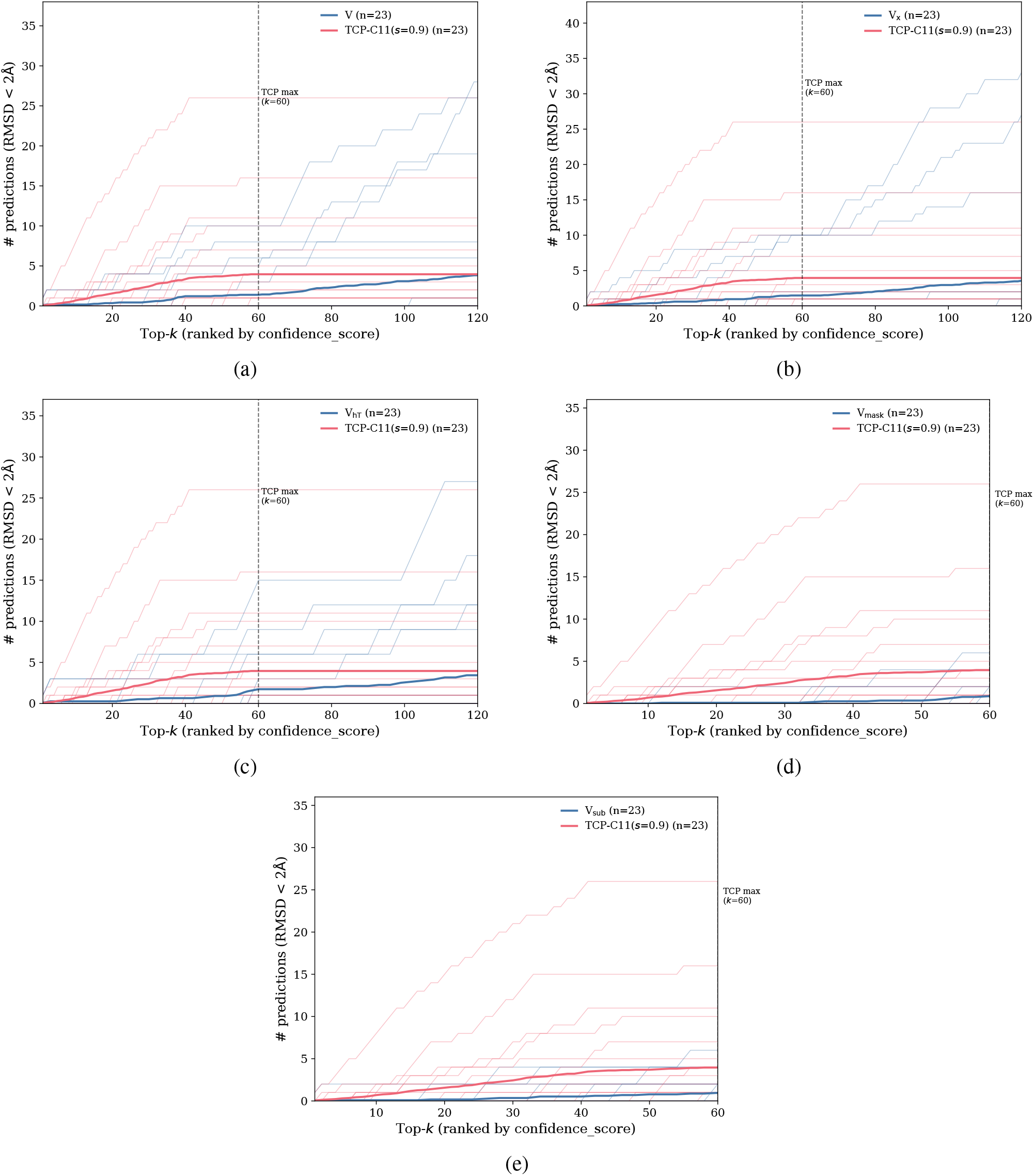
Top-*k* prediction quality (RMSD *<* 2 Å) ranked by confidence score (`confidence_score`). The y-axis represents the number of predictions with RMSD *<* 2 Å when we rank poses by confidence score and choose the top-*k* (x-axis) poses. Thin lines show individual targets; bold lines show the mean across *n* = 23 targets. The dashed vertical line marks the TCP budget (*k*=60), beyond which perturbation curves flatten. Only targets with at least one oracle-successful pose are included (*n* = 23 out of 57). Panels (a)–(e) compare TCP-C11 against baseline variants *V* (vanilla Boltz-2), *V*_*x*_ (Boltz-2x with steering potentials), *V*_hT_ (Boltz-2 with high diffusion temperature), *V*_mask_ (Boltz-2 with random maksing of MSA columns at rate 0.1), and *V*_sub_ (Boltz-2 with MSA depth reduced to 4086 rows), respectively. Since most targets have zero predictions below 2 Å, the mean flattens. Maximum prediction counts per method correspond to the *N* column in Table 2.

**Figure S9.**
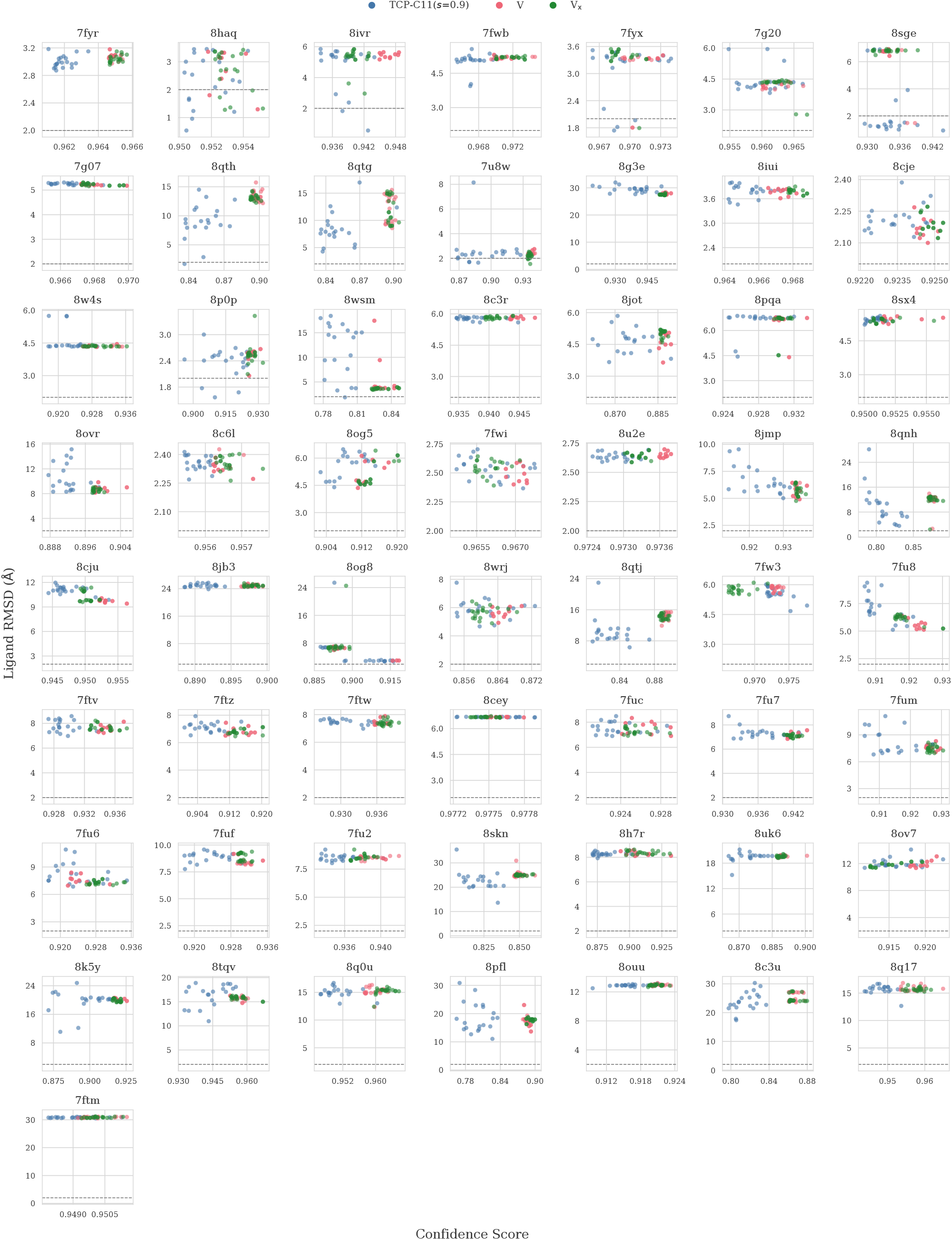
Full version of Figure S6.

**Figure S10.**
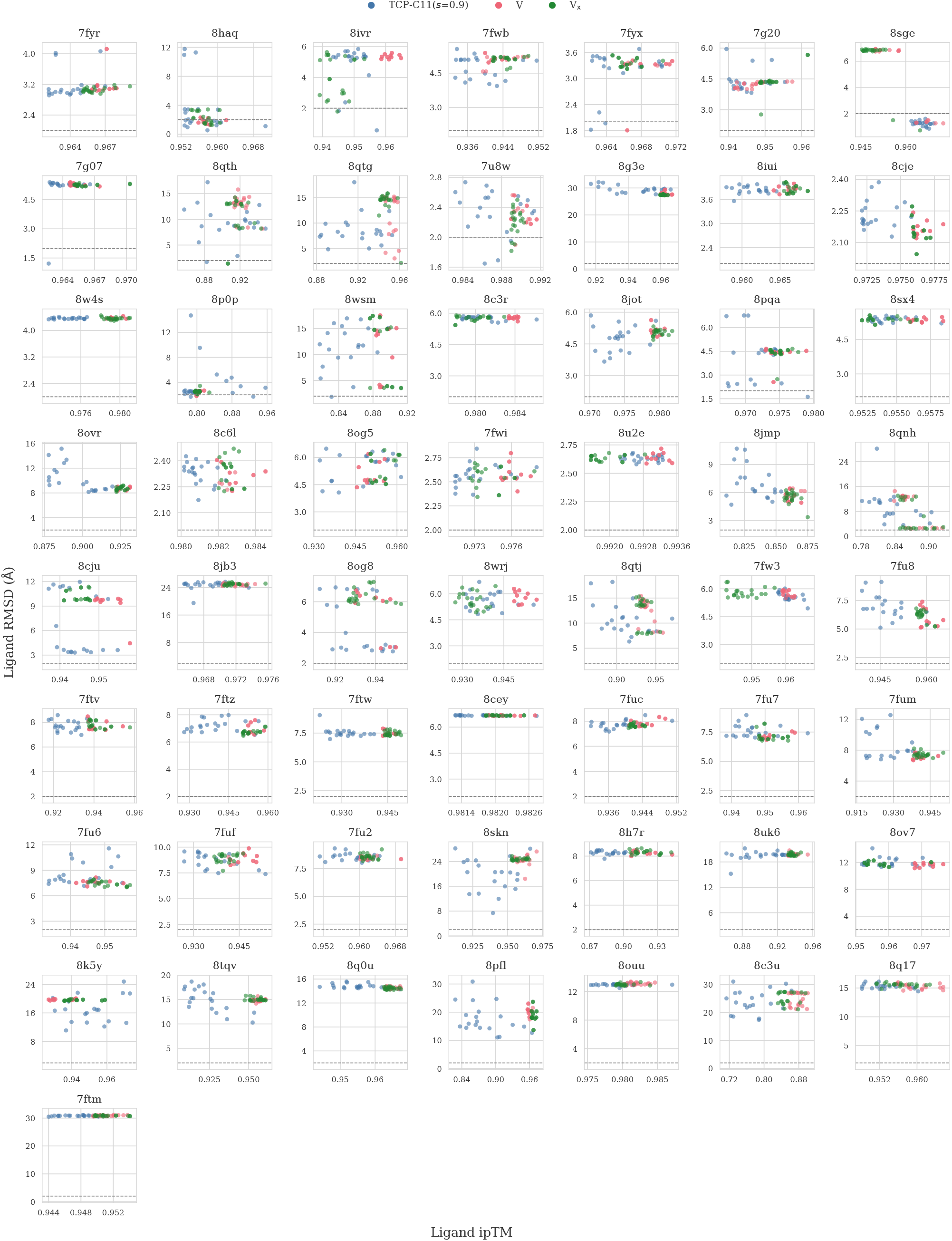
Full version of Figure S7.

## Notes

### Competing Interest Statement

The authors have declared no competing interest.

